# Sex Differences in *Drosophila* Intestinal Metabolism Contribute to Sexually Dimorphic Infection Outcome and Alter Gut Pathogen Virulence

**DOI:** 10.1101/2025.05.22.655590

**Authors:** Marko Rubinić, Aranzazu Arias-Rojas, Kaisy A. Martinez, Wioletta Klimek, Nicole Paczia, Kathirvel Alagesan, David Duneau, Igor Iatsenko

**Affiliations:** Research group Genetics of host-microbe interactions, Max Planck Institute for Infection Biology, Charitéplatz 1, 10117 Berlin, Germany; Humboldt-Universität zu Berlin, Faculty of Life Sciences, 10099 Berlin, Germany; Core Facility for Metabolomics and Small Molecule Mass Spectrometry, Max Planck Institute for Terrestrial Microbiology, Marburg, Germany; Max Planck Unit for the Science of Pathogens, Charitéplatz 1, 10117 Berlin, Germany; Centre for Cardiovascular Science, Queen’s Medical Research Institute, University of Edinburgh, Edinburgh, UK; Center for Ecology, Evolution and Environmental Changes (cE3c) & Global Change and Sustainability Institute (CHANGE), Faculty of Sciences, University of Lisbon (FCUL), Lisbon, Portugal; Broad Institute of MIT and Harvard, Cambridge, MA, USA; University of Texas at Arlington, 701 S Nedderman Dr, Arlington, TX 76019; Technische Universität Berlin, Straße des 17. Juni 135, 10623 Berlin

**Author notes:** Correspondence (I.I.).

**Keywords:** Gut Infection, Sexual Dimorphism, Immunometabolism, *Drosophila melanogaster*, *Pseudomonas*, Hfq

## Abstract

Sexual dimorphism in infection outcomes is a pervasive phenomenon, the underlying mechanisms of which remain incompletely understood. Here, utilizing *Pseudomonas entomophila* intestinal infection in *Drosophila,* we demonstrated that sex differences in intestinal redox processes contribute to female bias in susceptibility to gut infection. Female inability to overcome excessive pathogen-induced oxidative stress results in defecation blockage, pathogen persistence, and host death. Male flies exhibit increased carbohydrate metabolism and pentose phosphate pathway activity – a key antioxidant defense system. This allows males to withstand oxidative stress-induced defecation blockage and clear the pathogen from the intestine, resulting in survival. Additionally, *P. entomophila* showed increased expression of several virulence factors, including RNA-binding protein Hfq, in the female gut, contributing to female-biased virulence of *P. entomophila*. Thus, the effect of the gut metabolic environment on host defenses and pathogen virulence determines the sex differences in intestinal infection outcomes.

**Highlights:**

Intestinal transit of gut pathogen contributes to sexually dimorphic susceptibility to *Drosophila* gut infection.
Male bias in PPP favors pathogen clearance and recovery post-infection.
*P. entomophila* reacts differently to female gut environment, where higher levels of Hfq might contribute to virulence/lethality.

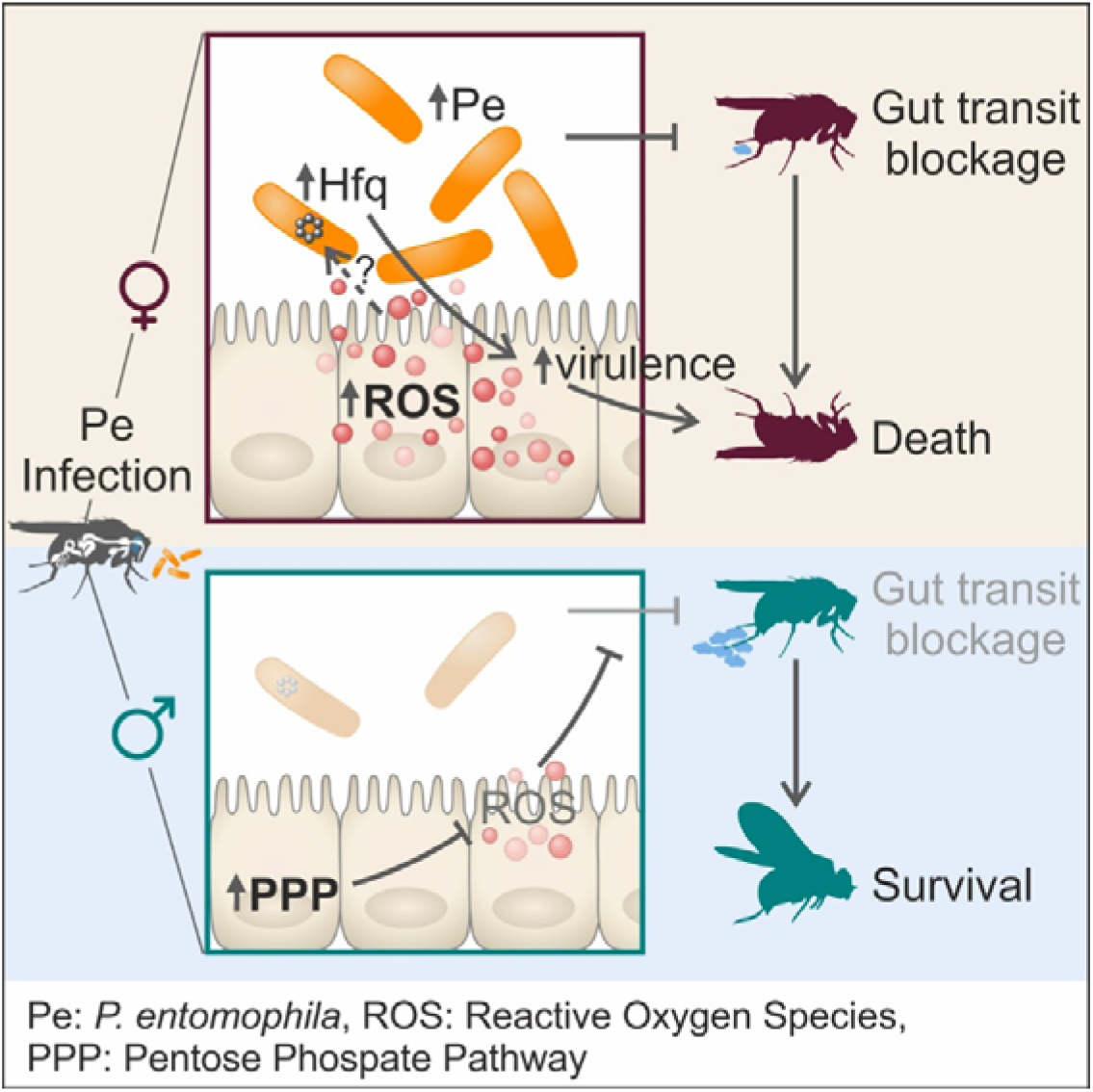

## INTRODUCTION

The sex is a determinant parameter of phenotypic and physiological heterogeneity of the organism^1–4^. Sex differences in immune defenses, response to infections, and anti-infective treatments have been often reported^5–11^. However, the mechanisms underlying these differences remain mostly unresolved, despite an emerging interest in understanding sexual dimorphism in immunity across taxa. Such lack of knowledge is not surprising, as it stems from the historical neglect of sex differences and the prevalent practice of utilizing only one sex in research studies. Understanding the complex interplay between sex and immunity has been further complicated by the fact that the dimorphism is often pathogen-specific, can be affected by the environment^12–16^, and depends not only on the host but also of the sex-specific pathogen response^17–19^.

Sexual dimorphism in response to gut infections remains particularly understudied, despite the accumulating evidence that certain gastrointestinal pathogens infect the host in sex-biased manner^11,13^. For instance, men are more frequently infected by *Campylobacter jejuni, Helicobacter pylori,* and *Vibrio spp.,* while enterohemorrhagic *Escherichia coli* infections are biased towards women^20,21^. Sexual dimorphism in some gastrointestinal infections could be attributed to immune differences between sexes^8,10^ or behavioral distinctions in hygiene and dietary practices^20^. Furthermore, the composition of gut microbiome was identified as another important factor that differs between sexes and influences immune system function^22,23^. The intricate interplay among immune responses, microbiota, behavior, environment, and genetic factors ultimately shapes the sex bias observed in the intestinal response to infections. Given the complexity of these interactions, it is not surprising that a comprehensive mechanistic understanding of sexual dimorphism in the outcomes of intestinal infections remains elusive. Experiments with genetically-tractable model organisms under controlled lab conditions represent a valuable approach to tackle the complex interplay between sex and infection outcome.

*Drosophila melanogaster* is a powerful model to study host-microbe interactions and host defense mechanisms due to a wide array of genetic tools that allow for the fine manipulation of cells and tissues both spatially and temporally^24–26^. Considering the increasing appreciation of sexually dimorphic physiology of fruit flies, they gain attention as a model to understand sexual dimorphism in immunity and infection outcome^4,5,15,27–33^. The advantages of *Drosophila* as a model for studying immune sex dimorphism lie in its genetic tractability, conserved immune pathways, well-defined microbiota, the ability to perform high-throughput screens and Genome-Wide Association Studies (GWAS). Additionally, the evolutionary conservation of key defense mechanisms in fruit flies^24,25^ allows for extrapolation of findings to other organisms, highlighting the broader relevance of insights gained from *Drosophila* studies on immune sex dimorphism.

Fruit flies rely on several defense mechanisms against intestinal pathogens^24,25^. The peritrophic matrix forms a physical barrier that shields epithelial cells from invading pathogens^34–36^. Additionally, the acidic region in the middle midgut represents a strong barrier, eliminating most of the ingested microbes via acid secretion^37–40^. The production of antimicrobial peptides (AMPs) and additional immune effectors, like iron-sequestration proteins, in specific regions of the gut represents an inducible arm of defense activated upon pathogen recognition^41–44^. While the Immune deficiency pathway (Imd) is a key regulator of antimicrobial response in the midgut of flies, subset of AMPs, like Drosomycin-like 2 (Drsl2) and Drosomycin-like 3 (Drsl3) are controlled by the JAK-STAT pathway^42^. The other major immune response pathway, the Toll pathway, functions specifically in the foregut and hindgut with less pronounce role in intestinal immunity^42^ but with a key role in sexual dimorphism to systemic infections^15^. Reactive oxygen species (ROS) are also rapidly produced in the gut by the Duox enzyme in the response to pathogen-secreted uracil^45^. While these ROS might be microbicidal against some microbes^46^, there is an accumulating evidence that Duox-produced ROS have signaling role in promoting gut peristalsis and clearance of the pathogens via defecation^47–49^. Duox was also shown to produce ROS in Malpighian tubules leading to the expression of cytokine Upd3 which stimulates epithelial renewal^50^. A number of additional signaling pathways are activated upon intestinal cell damage to initiate stem cell proliferation, tissue repair^42,51–53^ and resilience mechanisms^54^. Coordinated action of antimicrobial and tissue repair mechanisms is essential for fly survival to intestinal infection. While the described mechanisms were identified in females, we do not know whether and how they differ in males.

The intestinal defense mechanisms are very efficient in conferring the protection against transient microbes ingested with the food. Hence, there are only a few known microbes that can establish lethal infection in the *Drosophila* gut^55^. One of these microbes is *Pseudomonas entomophila* originally isolated from fruit flies^56^. *P. entomophila* can block intestinal defenses by producing toxins and proteases that degrade AMPs and compromise the integrity of peritrophic matrix^57–59^. During female *Drosophila* response to *P. entomophila*, excessive ROS accumulation results in oxidative stress, leading to translation blockage, nonreversible gut damage, and death of the majority of female flies^43,55–58,60^.

*P. entomophila* pathogenesis has not yet been described in male flies. Additionally, besides previously described differences between males and females in intestinal metabolic processes^61,62^, in the gut environment, there are a plethora of challenges, such as different pH levels, metabolite availability, or host immune effectors that gut pathogens need to react to^4,63^. Whether and how males and females differ in these processes and if these differences contribute to gut immune defenses is not known. Similarly, gut metabolic differences between sexes might distinctly affect pathogen virulence, thus contributing to sexual dimorphism in infection susceptibility^64^. This possibility remains to be tested.

We studied in-depth the sex difference in *Drosophila* susceptibility to an intestinal infection, using the *Drosophila* natural bacterial pathogen *P. entomophila*. We used transcriptomics, proteomics, GWAS, and targeted metabolomics to determine the mechanisms underlying the difference between sexes. We showed that male bias in basal gut NADPH-producing enzyme levels contributes to the antioxidant response to gut infection, favoring regular intestinal transit, bacterial clearance, and overall better survival than female flies. We undertook a dual proteomics approach and identified *P. entomophila* proteins differentially expressed in male and female gut environments. A RNA chaperone Hfq - a known regulator of virulence^65–67^ was the *P. entomophila* protein which was the most abundant in females guts compared to male guts, suggesting that the host environment regulates pathogen virulence. Overall, our findings reveal a mechanism by which carbohydrate metabolism shapes sexual dimorphism in susceptibility to infection by affecting ROS’s effects on the host and the pathogen.

## RESULTS

### *Drosophila* susceptibility to gut infection is sexually dimorphic

To study to which extent *Drosophila* is sexually dimorphic in susceptibility to intestinal infection, we infected male and female flies of the same genetic background by feeding them with a mix of a pathogen suspension and sucrose using a previously established protocol^43^ (Figure 1A). We used natural *Drosophila* pathogen *P. entomophila* that was shown to be highly lethal to female flies^43,56^. Notably, we observed that the vast majority of male flies of three commonly used *Drosophila* lab strains; *w*^1118^ iso, Canton S, and Oregon R, survived *P. entomophila* infection, while 70 to 80% of female flies were killed (Figure 1B, 1B’), demonstrating that sexual dimorphism in susceptibility to intestinal infection is common among lab ‘wild-type’ genotypes. To further demonstrate the generality of sexual dimorphism in susceptibility to gut infection among genetically distinct individuals, we tested survival post-exposure to *P. entomophila* in environment-controlled conditions (see: Methods) of 183 *Drosophila* melanogaster genetic reference panel (DGRP) lines^68^ allowing to test the proportion of genotypes with a given proportion of dimorphism. As anticipated from a previous study^60^, we observed variation among DGRP lines in susceptibility to *P. entomophila* in both sexes (Figure 1C, S1A, S1B). Correlation between hazard ratios of both sexes shows a high ratio of positive linear relationships between the sex and survival outcomes following an infection (Figure 1C). Since we tested male and female flies of each DGRP line in parallel, we could calculate the sexual dimorphism in survival for each line (Figure 1D). Among the 120 lines that showed statistically significant sexually dimorphic susceptibility, 87.5% (105/120) exhibited phenotype observed with lab strains - *P. entomophila* being more lethal to female files. Collectively, these results confirm the generality of sex differences in susceptibility to intestinal infection among genetically distinct hosts.

**Figure 1.**
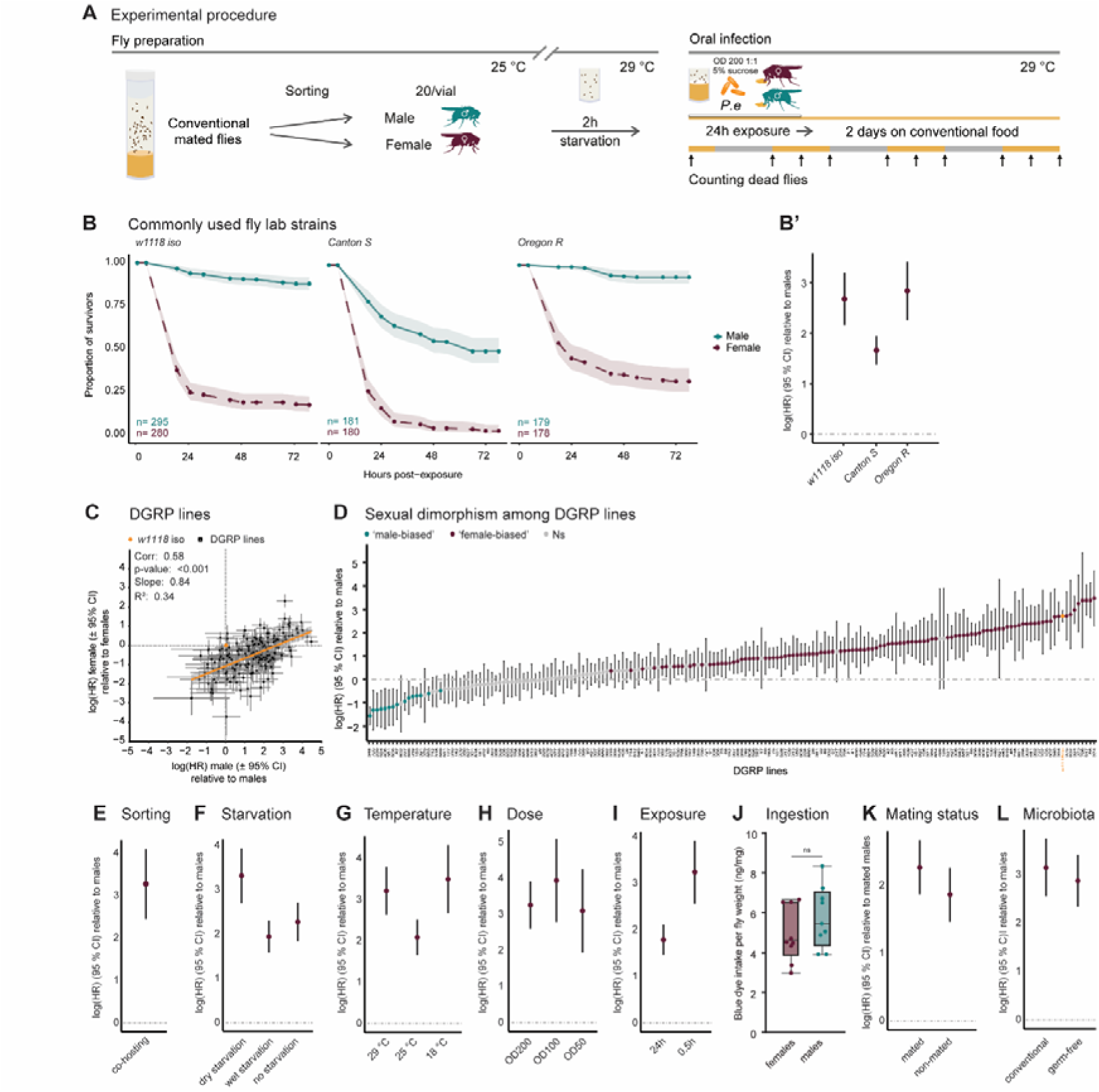
*Drosophila* susceptibility to gut infection is sexually dimorphic. **(A)** Graphical overview of experimental procedure to investigate sex differences in survival to intestinal infection. **(B and B’)** Survival curves with 95% confidence intervals (shaded area) and hazard ratios with 95% confidence intervals of commonly used lab strains (*w*^1118^ iso, Canton S, Oregon R) upon exposure to *P. entomophila*. **(C)** A scatter plot representing correlation in the variation of hazard ratios of female and male flies (each relative to the appropriate sex of the *w^1118^* iso strain) within the same DGRP line. **(D)** Variability in sexual dimorphism in susceptibility to *P. entomophila* infection across 183 DGRP lines (DGRP lines where female flies were more susceptible compared to male flies are highlighted in plum, those where male flies were more susceptible compared to female flies are highlighted in teal. The reference *w^1118^* iso strain is in orange despite the female’s bias). **(E – I)** Hazard ratios with 95% confidence intervals of *w^1118^* iso upon exposure to *P. entomophila* under different protocol conditions; **(E)** co-hosting, **(F)** dry starvation, wet starvation, and no starvation, **(G)** 29 °C, 25 °C, and 18 °C, **(H)** OD200, OD100, and OD50, **(I)** 24 h and 0.5 h. **(J)** Intake of blue dye mixed with *P. entomophila* during 0.5 h of exposure of female and male *w^1118^* iso flies. Each dot represents a pool of five flies taken from different infection vials, and the experiment was independently performed on three different days. **(K)** Hazard ratios with 95% confidence intervals of mated and non-mated *w^1118^* iso females upon exposure to *P. entomophila* compared to mated *w^1118^* iso males. **(L)** Hazard ratios with 95% confidence intervals of conventional and germ-free *w^1118^* iso females upon exposure to *P. entomophila* compared to appropriate male flies. Throughout the panels, error bars represent 95% confidence intervals (CIs). Non-overlapping CIs between treatments or genotypes indicate statistically significant differences (p < 0.05). Similarly, when a 95% CI does not overlap with the dashed reference lines, it indicates a significant difference from the reference value (generally female). For detailed sample sizes and statistical analyses, see Table S9.

Environmental factors can differently affect each sex and, consequently, susceptibility to infection. Therefore, we tested how stable sexual dimorphism in susceptibility to *P. entomophila* infection is under different protocol conditions. Sexual dimorphism was equally pronounced when male and female flies were infected and co-hosted in the same vial and when they were infected and separated in different vials (Figure 1E, S1C). Thus, sex differences in survival are not due to potential variability between vials in infectivity or due to the effect of isolation. Since typical oral infection protocol includes a 2 h starvation step in an empty vial (dry starvation), which might negatively affect females more than males, we tested alternative starvation strategies. We still observed increased susceptibility of female flies to infection following wet starvation (vials with 1% agar) or without starvation (Figure 1F, S1D). We also found that dimorphism was still striking at different temperatures, although stronger at our two extremes (Figure 1G, S1E). These results indicate that the sexual dimorphism was robust to various environmental conditions. Furthermore, differences in gut infection can occur due to differences in feeding behavior that were previously reported between sexes^69,70^. To test this, we took two different approaches: comparing sexual dimorphism using a lower dose of *P. entomophila* and exposing flies to *P. entomophila* for 0.5 h (referred to as ‘0.5 h protocol) to ensure that the potential difference in continuous ingestion does not explain the sexual dimorphism. Differences between males and females in survival were present when different doses of *P. entomophila* were used (Figure 1H, S1F) and in the 0.5 h protocol (Figure 1I, S1G). Finally, we confirmed that initial pathogen ingestion in the 0.5 h protocol does not statistically differ between sexes (Figure 1J). This suggests that the difference in susceptibility is not due to a difference in initial pathogen intake. Given a known trade-off between reproduction and immunity^71^, we tested whether females might be more susceptible to infection due to higher investment in reproduction at the expense of immunity. As expected, virgin females, compared to mated ones, survived *P. entomophila* infection better (Figure 1H, S1H’). However, they were still more susceptible than males (Figure 1K, S1H). Thus, the increased susceptibility of females to infection is not due to the immunosuppressive effect of reproduction. Susceptibility to gut infection can also be affected by the microbiota composition^39,72,73^. However, germ-free flies showed sex differences comparable to conventional flies in susceptibility to infection (Figure 1L, S1I), excluding the possibility that sexual dimorphism is determined by microbiota.

Overall, sexual dimorphism in susceptibility to *P. entomophila* intestinal infection is pervasive among host genetic backgrounds and is independent of host reproductive status, feeding behavior, microbiome differences or common environmental factors, warranting further exploration of the underlying mechanisms.

### Previously described defense pathways are not the leading cause of sex differences in susceptibility

Since intestinal immune defenses have not yet been characterized in male flies, we investigated the transcriptional response to *P. entomophila* infection of male alongside of female flies. We performed bulk RNA-seq of dissected guts of infected male or female *w*^1118^ iso flies at 6 h and 16 h time points after exposure to *P. entomophila*. Differential expression analysis between pathogen-exposed and non-exposed controls was performed within each sex (Figure 2A, S2A, Table S1). We observed a lower number of significantly upregulated genes in males (6 h: 837 in males vs 1424 in females, 16 h: 1414 males vs females 1564). However, 81% (678 out of 837) of all genes induced in males at 6 h post-infection was also upregulated in females (see Table S1 for a list of overlapping and unique genes). Although the overlap was lower for 16 h (64% (913 out of 1564)), these results show overlay in the transcriptional response of males and females to *P. entomophila* infection. Specifically, we found that males and females share upregulated genes involved in antimicrobial defense (AMPs, Tsf1), stress response (Turandots, Hsps, Gstds), stem cell activation and epithelial renewal (EGFR and JAK-STAT pathways), gut structure (Cry, peritrophin) (Figure 2A, 2C, S2A, Table S1). Genes encoding digestive enzymes were repressed after infection in both sexes. These results are consistent with a previous study that analyzed the female response to *P. entomophila* infection^43^ and suggest that sexual dimorphism in survival is unlikely primarily due to sex differences in these processes.

**Figure 2.**
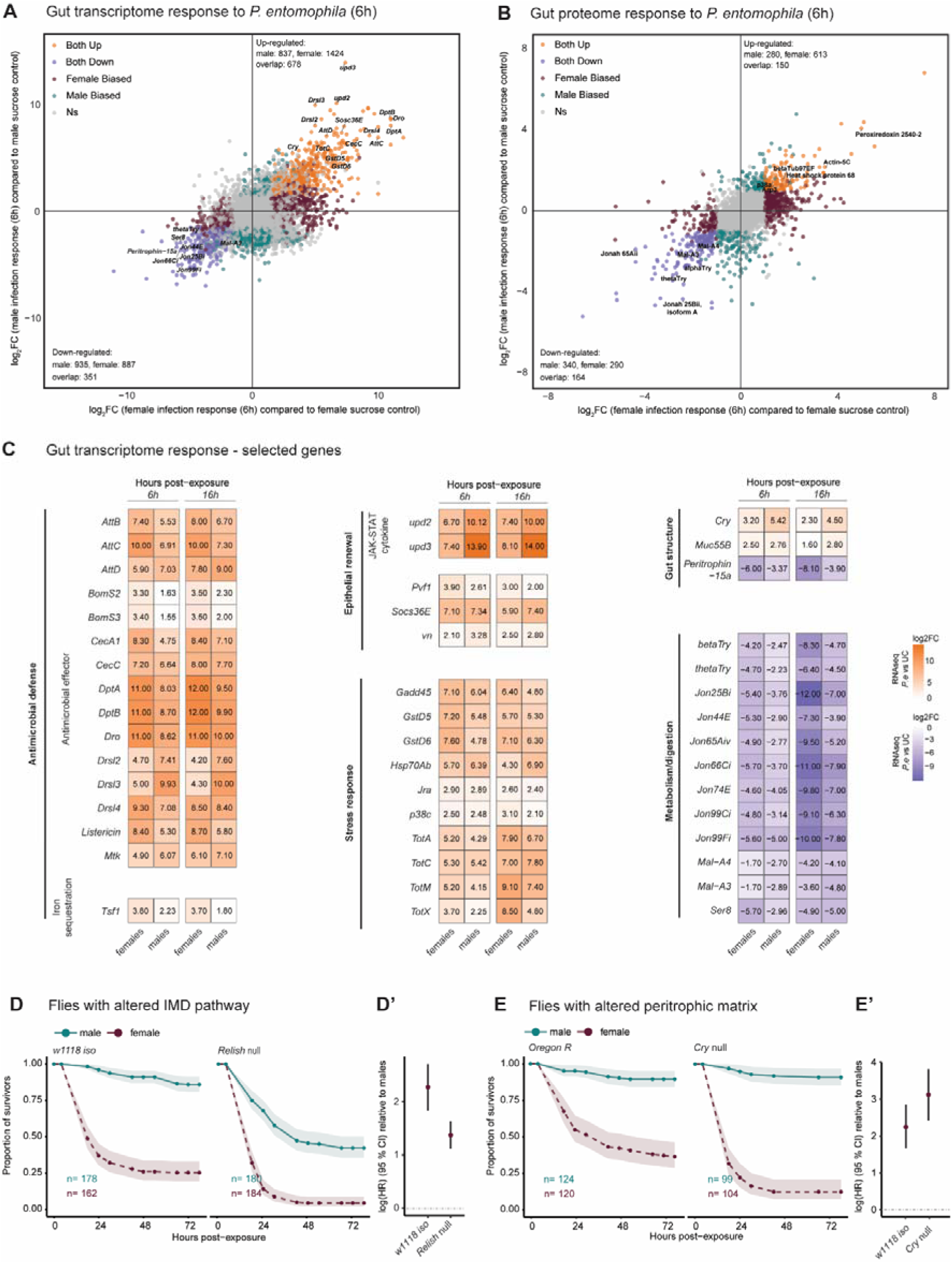
Previously described defense pathways are not the leading cause of sex differences in susceptibility. **(A and B)** Scatter plots representing log_2_FC of **(A)** gene expression or **(B)** protein abundance in female (x) vs male (y) guts 6 h post-exposure to *P. entomophila* (infected guts compared to sucrose-fed (control) guts of matching sex). Significance cut-offs: **(A)** padj < 0.1, |log_2_FC| > 1.5, and **(B)** padj < 0.05, |log_2_FC| > 1. **(A)** (N = 3 independent samples, each with 30 pooled guts). **(B)** (N = 5 independent samples, each with 30 pooled guts). **(C)** Heatmaps showing differences (log_2_FC, infected guts compared to control guts) in the expression of selected genes of each sex at 6 h and 16 h post-exposure to *P. entomophila*. **(D and D’)** Survival curves with 95% confidence intervals (shaded area) and hazard ratios with 95% confidence intervals of *w^1118^* iso (control) and *Relish^E20^*loss-of-function mutant upon exposure to *P. entomophila*. **(E and E’)** Survival curves with 95% confidence intervals (shaded area) and hazard ratios with 95% confidence intervals of *Oregon R* (control) and *Cry* loss-of-function mutant upon exposure to *P. entomophila*. For detailed sample sizes and statistical analyses, see Table S9.

Since *P. entomophila* causes translation blockage in female flies^43^, we additionally undertook a proteomics approach under the same conditions as RNAseq to look for potential differences at the proteome level. Similarly to RNAseq, we observed a higher number of upregulated proteins in female flies (6 h: 278 in males vs 611 in females, 16 h: 359 males vs females 911). Consistent with transcriptomics, we detected substantial overlap in the proteome response of males and females to *P. entomophila* infection (6 h: 53.6% (149/278), 16 h: 79.7% (296/359), see Table S2 for a list of proteins). Among proteins that were upregulated in both sexes, we detected stress response proteins (heat shock proteins, p38a, peroxiredoxin 2540), proteins with a role in the cytoskeleton and epithelial renewal (betaTub56D, betaTub97EF, Tubulin beta-3 chain, Actin-related protein 3, Actin-5C) (Figure 2B, S2B, S2C, Table S2). Repressed proteins were enriched with digestive enzymes (trypsins, proteases, maltases, glucosidases). Notably, proteomic analysis in both sexes has not detected any immune effectors, like AMPs, being induced by *P. entomophila* infection. This might be a consequence of translational blockage by *P. entomophila,* as numerous AMPs were transcriptionally induced by infection. Next, we decided to conclusively test some of the defense pathways using respective mutants.

Considering the important role of the IMD pathway in the defense against *P. entomophila* gut infection^59^ and that the IMD pathway was upregulated in both sexes (Figure 2A, 2C, S2A), we tested the contribution of this pathway to sexual dimorphism. Although *Relish* mutants, deficient in the IMD pathway, were more susceptible to *P. entomophila* infection, they still exhibited sexual dimorphism. Thus, the IMD pathway is important in defense against *P. entomophila,* however, it is not sufficient to explain the observed sexual dimorphism (Figure 2D and 2D’). Since the peritrophic matrix was described as an important defense barrier against *P. entomophila* infection in female flies^34,35^, we tested whether sex differences in this physical barrier could explain the differences in survival. We infected *Drosocrystallin* (here referred to as *Cry*) loss-of-function mutant deficient in the peritrophic matrix and still observed sexual dimorphism (Figure 2E and 2E’). Lastly, considering that the Toll pathway has been suggested to have a major role in sexual dimorphism in *Drosophila* systemic immunity among various bacteria^15^, we tested whether sexual dimorphism in the Toll pathway is connected to the sex difference in survival after *P. entomophila* infection. We tested two loss-of-function mutants in Toll pathway: *spätzle* (*spz*) and *modular serine protease* (*modsp*). Both mutants exhibited sex differences in survival comparable to wild-type flies (Figure 2D, S2D’, S2E, S2E’), indicating that the Toll pathway does not underlie the observed sexual dimorphism.

Overall, we described and compared the response of female and male flies to ingested *P. entomophila* on transcriptomic and proteomic levels that leave opportunities for further exploration of potential candidates with a role in intestinal defense. Considering that previously described defense mechanisms do not explain the sexual dimorphism in susceptibility to *P. entomophila*, we further explored the underlying mechanisms.

### *P. entomophila* causes sex-biased defecation blockage

*Drosophila* survivorship following infection is determined by the interplay of immune activity and pathogen burden at critical time points^74^. Therefore, we measured *P. entomophila* load in infected flies to assess whether males and females differ in the ability to control pathogens. To reflect only the behavior of bacteria in the gut and not the influence of possible reingestion of bacteria, bacterial load was measured using flies exposed to *P. entomophila* for 0.5 h (which has a similar effect on survival as 24 h infections (Figure 1I, S1G)). We tested pathogen burden at different time points by homogenizing five pooled flies, plating on Luria Broth (LB) plates, and counting colony-forming units (CFU). While the *P. entomophila* burden remained high through different time points, as previously described in female flies^57^, male flies started clearing *P. entomophila* already at early time points (2 h), and completed pathogen clearance approximately 6 h post-exposure (Figure 3A).

**Figure 3.**
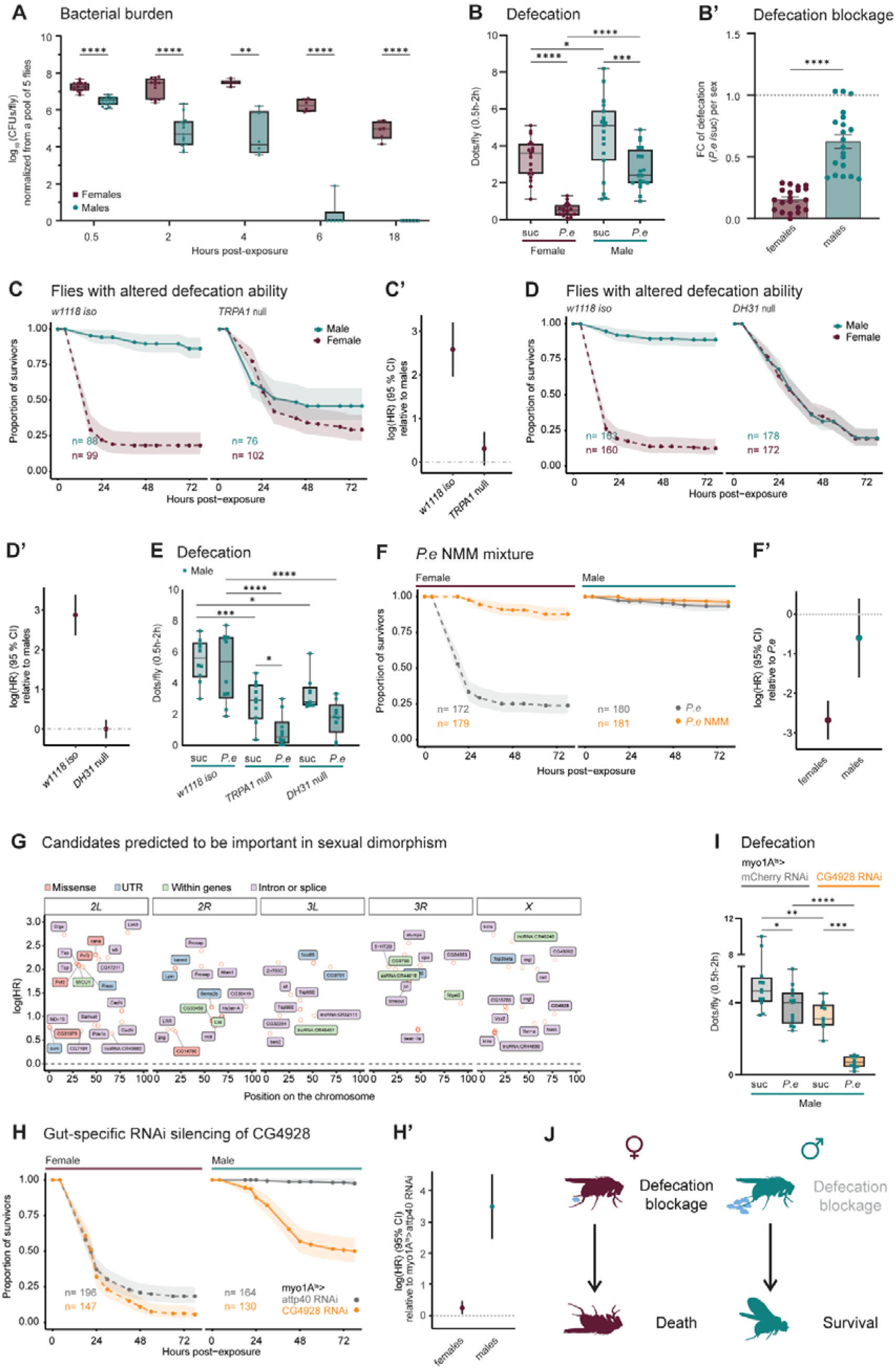
*P. entomophila* causes sex-biased defecation blockage. **(A)** Pathogen abundance (CFUs) at 0.5, 2, 4, 6, and 16 h post-exposure to *P. entomophila* in 0.5 h protocol. One dot indicates biological replicate (pool of five flies). Significance by Mixed-effects model (REML) with Šídák’s multiple comparisons test for each Time point. **(B and B’)** The defecation rate of female and male *w^1118^* iso flies measured 0.5 to 2 h after exposure to blue-dyed sucrose (control) or *P. entomophila* in 0.5 h protocol. **(B)** Significance by two-way ANOVA with Šídák’s multiple comparisons test. Interaction (Sex x Treatment) p = n.s. (**B’)** Fold change calculated from mean values of sucrose control of appropriate sex from the same biological repeat. Significance by Mann Whitney test. **(C – D’)** Survival curves with 95% confidence intervals (shaded area) and hazard ratios with 95% confidence intervals of *w^1118^* iso (control) and two mutants with impaired defecation ability: **(C – C’)** *TRPA1* loss-of-function mutant and **(D – D’)** *DH31* loss-of-function mutant. **(E)** The defecation rate of male *w^1118^* iso (control) flies and two mutants with impaired defecation ability (*TRPA1* loss-of-function mutant and *DH31* loss-of-function mutant) measured 0.5 to 2 h after exposure to blue-dyed sucrose (control) or *P. entomophila* in 0.5 h protocol. Significance by ordinary one-way ANOVA, Šídák’s multiple comparisons test. **(F and F’)** Survival curves with 95% confidence intervals (shaded area) and hazard ratios with 95% confidence intervals of *w^1118^* iso flies upon exposure to *P. entomophila* and *P. entomophila*/N-methyl maleimide mixture. **(G)** Candidate genes underlying natural variation in the sexual dimorphism to *P. entomophila* gut infection. **(H and H’)** Survival curves with 95% confidence intervals (shaded area) and hazard ratios with 95% confidence intervals of *myo1A^ts^>attp40* RNAi (control) and *myo1A^ts^>CG4928* RNAi upon exposure to *P. entomophila*. **(I)** The defecation rate of male *myo1A^ts^>attp40* RNAi (control) and *myo1A^ts^>CG4928* RNAi flies measured 0.5 to 2 h after exposure to blue-dyed sucrose (control) or *P. entomophila* in 0.5 h protocol. Significance by two-way ANOVA with Šídák’s multiple comparisons test. Interaction (Genotype x Treatment) p = n.s. **(J)** Graphical summary illustrating the link between sexual dimorphism in defecation and survival. Throughout the panels, boxplots and dot plots show median and interquartile ranges (IQR); whiskers show the full data range. For detailed sample sizes and statistical analyses, see Table S9.

Next, we investigated how males clear the pathogen: via killing in the intestine or expulsion through gut peristalsis. To test the previously described mechanism of pathogen clearance via expulsion from the intestine by gut peristalsis, we infected flies with a mixture of *P. entomophila* and blue color dye that allowed to estimate the intestinal excretion by scoring the number of defecation spots deposited within 1.5 h after a 0.5 h exposure^48^. We found that *P. entomophila* significantly reduced defecation in both males and females as compared to sucrose controls (Figure 3B). However, the effect was much stronger in females, leading to almost complete defecation blockage (Figure 3B’) and, thus, likely preventing the pathogen clearance via expulsion. We genetically and chemically manipulated intestinal transit during *P. entomophila* infection to test the causality between defecation and survivorship. Upon bacterial ingestion, the gut produces ROS sensed by evolutionarily conserved Transient Receptor Potential A1 channel (TRPA1) in enteroendocrine cells that, as a result, release Diuretic Hormone 31 (DH31), which activates gut contraction favoring pathogen expulsion^48,49^. When we compared previously described loss-of-function mutants lacking *TRPA1* or *DH31*, the sexual dimorphism in susceptibility was not present anymore; males were as susceptible as females to *P. entomophila* (Figure 3C, 3C’, 3D, 3D’). Since *TRPA1* and *DH31* mutant males exhibited increased susceptibility to infection as compared to control males, we measured defecation in males to test whether higher susceptibility of mutants is linked to reduced ability to expel the pathogen. As anticipated, males from both mutants had reduced defecation in comparison to wild-type control (Figure 3E). We then tested whether increasing the intestinal transit decreased susceptibility by exposing infected flies of both sexes with N-methyl maleimide (NMM), a chemical that increases the intestinal transit by activating TRPA1^48^. The low susceptibility of males was not affected. However, it reduced strongly the female susceptibility, bringing them close to those of males (Figure 3F, F’). Combined, we showed that pathogen causes defecation blockage in infected flies in a sexually dimorphic manner leading to differential susceptibility of male and female flies to the same infection.

We investigated the genetic basis of the susceptibility to *P. entomophila* oral infection in each sex (Figure 1A, S1B), and of their dimorphism (Figure 3G) using a GWAS. To do so, we used the genetic and phenotypic variations present in each sex in the DGRP lines and in *w*^1118^ iso, the latter being used as a reference across the experiments. The candidate SNPs associated with male susceptibility, female susceptibility and sexual dimorphism in susceptibility are listed in Table S3 and curated in Figure 3G, S3A, S3B based on the functional impact of the mutation (i.e. missense, UTR, within gene or intron/splice region), the p value, and the size of the effects observed on the DGRP lines (i.e. log (hazard ratio).

Then, to validate the potential contribution of those SNPs to variation in susceptibility to *P. entomophila*, we decided to test whether the change in expression of genes associated with those SNPs will result in a difference in susceptibility. However, SNP variation has consequences on the whole body and many of our candidates are expected to be linked to their impact on other tissues or on gut development. We were interested to find those candidates affecting the processes during the infection, therefore we targeted the expression of some candidate genes (listed in Table S3) specifically in enterocytes at the adult stage. The strongest effect on sexual dimorphism in survival was observed when we reduced enterocyte expression of *CG4928* (Figure 3H, 3H’, S3C, S3C’). *CG4928* is an understudied transporter predicted to enable potassium channel regulator activity and transmembrane transport of potassium^75^. Considering that *CG4928* shows high expression level in the gut^76^ and known link between potassium and intestinal motility^77,78^, we hypothesized that *CG4928* affects susceptibility to infection via altering pathogen clearance by defecation. Indeed, we found that while control males still retained defecation ability after infection, in *CG4928* RNAi male’s defecation was almost completely inhibited (Figure 3I). Hence, our GWAS approach provided an additional unbiased confirmation of the link between survival and defecation rates. Additionally, these results link *CG4928* with defecation ability in *Drosophila* (even in non-infected sucrose-fed conditions) possibly via affecting gut contractions via potassium disbalance, which remains to be explored.

Altogether, these results support a scenario where males, due to their reduced susceptibility to *P. entomophila*-induced defecation blockage, can efficiently clear the pathogen via gut peristalsis and thus survive the infection (Figure 3J).

### Male gut antioxidant capacity contributes to the attenuated impact of *P. entomophila* on defecation blockage

Considering the known role of ROS in gut peristalsis^47–49^, we investigated whether male and female differences in ROS production or sensitivity contribute to the observed differences in survival and pathogen eviction via defecation. Given that in female flies*, P. entomophila* induces a high level of intestinal ROS^43,60^, which suppresses immune and repair processes via translational blockage^43^, we hypothesized that males might prevent or resist such excessive oxidative stress and thus survive the infection. To compare oxidative stress levels during infection in male and female flies, we measured 2’,7’-dichlorofluorescein (H_2_DCF), which reflects general oxidative stress levels^79^. Consistent with previous studies^43,60^, we detected a significant increase in oxidative stress in female guts after *P. entomophila* ingestion (Figure 4A). Such an increase was not observed in male flies (Figure 4A). Importantly, Sleiman et al. ^60^ showed that DGRP lines that are less susceptible to *P. entomophila* infection have low ROS levels after infection when compared to sensitive lines. Because of this correlation between oxidative stress and susceptibility to *P. entomophila* infection, we wondered whether male flies have higher antioxidant capacity, which can be tested by exposing flies to compounds that induce oxidative stress in flies, such as commonly used oxidizing agent paraquat^43^. Male flies were less susceptible to paraquat than females (Figure 4B, 4B’), indicating they can handle oxidative stress better.

**Figure 4.**
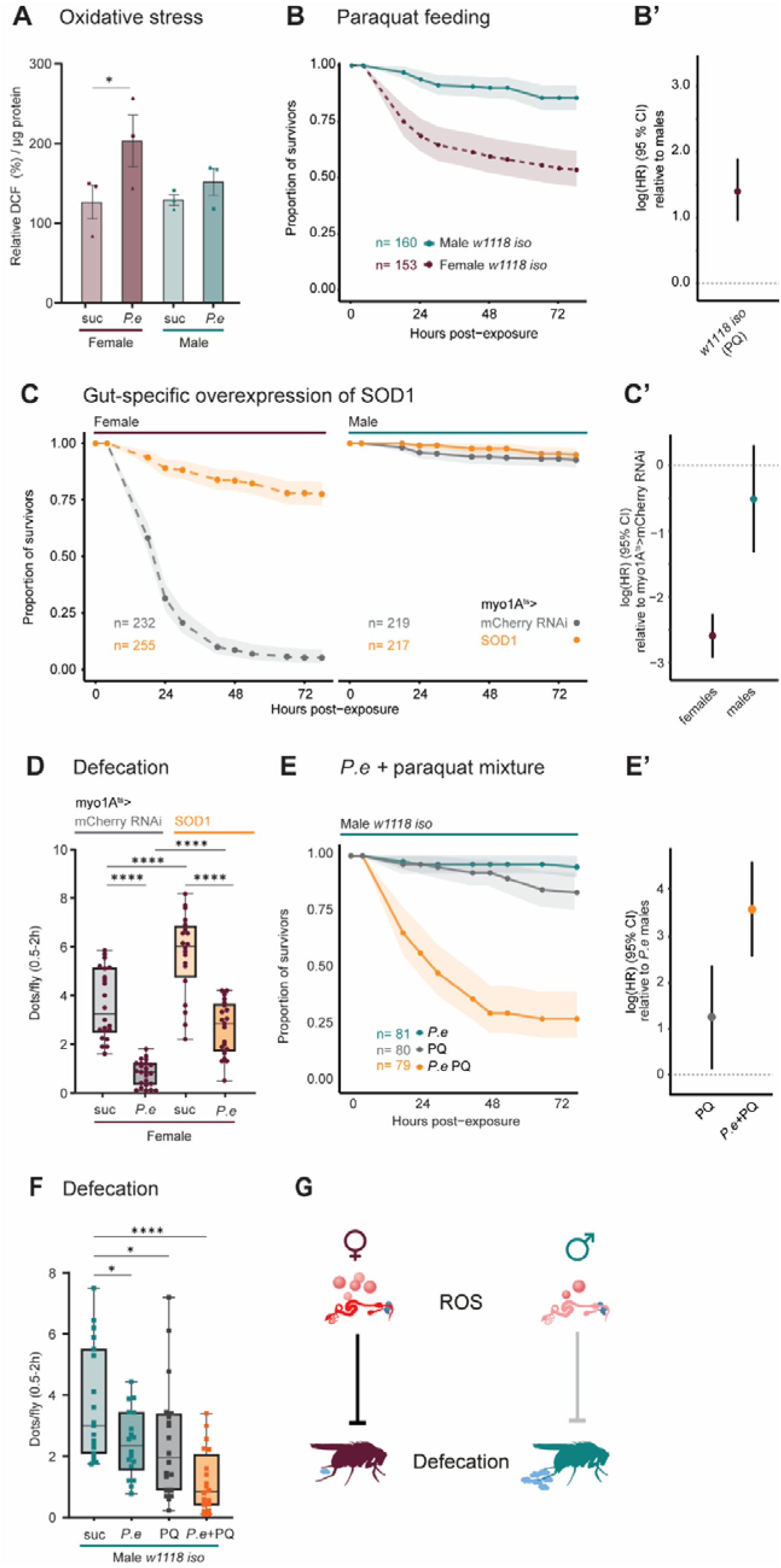
Male gut antioxidant capacity contributes to the attenuated impact of *P. entomophila* on defecation blockage. **(A)** Reactive oxygen species (ROS) measured as percent of 2′,7′-dichlorofluorescein (DCF) relative fluorescence units (RFU) normalized per homogenized gut samples (N = 3 independent samples represented by different shape, n = 30 guts per sample). Mean ± SE. ∗p < 0.05 by two-way ANOVA with Šídák’s multiple comparisons test. Interaction (Sex x Treatment) p = n.s. **(B and B’)** Survival curves with 95% confidence intervals (shaded area) and hazard ratios with 95% confidence intervals of female and male *w^1118^* iso flies upon exposure to paraquat (4 h protocol). **(C – C’)** Survival curves with 95% confidence intervals (shaded area) and hazard ratios with 95% confidence intervals of female and male *myo1A^ts^>mCherry* RNAi (control) and *myo1A^ts^>UAS-SOD1* (BL24750). **(D)** The defecation of female *myo1A^ts^>mCherry* RNAi (control) and *myo1A^ts^>UAS-SOD1* (BL24750) flies measured 0.5 to 2 h after exposure to blue-dyed sucrose (control) or *P. entomophila* in 0.5 h protocol. Significance by two-way ANOVA with Šídák’s multiple comparisons test. Interaction (Genotype x Treatment) p = n.s. **(E and E’)** Survival curves with 95% confidence intervals (shaded area) and hazard ratios with 95% confidence intervals of male *w^1118^* iso flies upon exposure to *P. entomophila*, paraquat, and *P. entomophila*/paraquat mixture. **(F)** The defecation of male *w^1118^* iso flies measured 0.5 to 2 h after exposure to blue-dyed sucrose (control), *P. entomophila*, paraquat, or *P. entomophila*/paraquat mixture in 0.5 h protocol. Significance by two-way ANOVA with Šídák’s multiple comparisons test. Interaction (Sex x Treatment) p = n.s. **(G)** Graphical summary illustrating the link between sexual dimorphism in intestinal infection-induced ROS levels and defecation. For detailed sample sizes and statistical analyses, see Table S9.

Then, we wondered whether sex differences in *P. entomophila-*induced oxidative stress contribute to observed differences in survival. We tested the effect of N-acetyl-L-cysteine (NAC), a commonly used antioxidant compound^80^. We observed that both infected female and male flies survived significantly longer when, after infection, they were fed NAC-supplemented sucrose as compared to sucrose alone (Figure 4A, S4A’), confirming that increased oxidative stress affects female susceptibility to *P. entomophila*. Additionally, we tested whether the expression of antioxidant enzymes in enterocytes affects fly susceptibility to infection. We observed an increase in survival of infected flies with overexpression of cytosolic antioxidant enzyme superoxide dismutase 1 (SOD1) with two distinct transgenic lines (Figure 4C, 4C’, S4B, S4B’) and a minimal effect of overexpression of mitochondrial SOD2 (Figure 4C, S4C’). Additionally, we fed SOD1-overexpressing female flies with *P. entomophila,* paraquat, or a mixture of *P. entomophila* and paraquat to test whether we can override the antioxidant capacity of SOD1 with paraquat. Although SOD1-overexpressing flies survived better on *P. entomophila* or paraquat alone compared to control flies, they could not survive the excessive oxidative stress caused by the *P. entomophila/*paraquat mixture (Figure 4D, S4D’). Overall, these results indicate that cytosolic (and not mitochondrial) oxidative stress is a determinant of susceptibility to infection in female flies, and differences between sexes in oxidative stress contribute to sexual dimorphism in susceptibility to *P. entomophila* gut infection.

To further broaden our understanding of the relationship between oxidative stress, susceptibility to infection, and newly described defecation blockage, we tested whether altering oxidative stress genetically or chemically will also affect the amount of defecation. Overexpression of SOD1 in female flies also significantly increased defecation after infection (Figure 4D), suggesting that reducing excessive ROS in females restores gut peristalsis and pathogen clearance. When we did the reciprocal experiment and increased oxidative stress in males by combining *P. entomophila* with paraquat, we observed not only high susceptibility of males to *P. entomophila* (mixed with paraquat) infection (Figure 4E, 4E’) but also significantly reduced defecation (Figure 4F), suggesting that the reduced ability to clear the pathogen affects survival. These findings show that excessive oxidative stress inhibits not only immune and repair processes, as previously shown^43^, but also blocks intestinal transit and prevents pathogen clearance. Males’ ability to prevent the infection-induced oxidative burst allows them to clear the pathogen via intestinal peristalsis and survive the infection (Figure 4G).

### Abolishing differences in NADPH metabolism removes sexual dimorphism in susceptibility to infection

Next, we wanted to understand the mechanisms that mediate males’ resistance to oxidative stress. The transcriptomic and proteomic analyses in both uninfected sexes (Table S4) showed male-biased upregulation of genes involved in carbohydrate metabolism. This was suggested by the functional enrichment analysis of transcripts in male intestines (Figure 5A), and is consistent with a previous study^62^. In particular, numerous transcripts encoding steps in glycolysis, the pentose phosphate pathway (PPP), and the tricarboxylic acid cycle were expressed stronger in males (Figure 5B). The proteomic analysis confirmed that male intestines have increased abundance of proteins involved in carbohydrate metabolism than females (Figure 5A, S5B). To illustrate that these higher levels result in a higher pathway activity in males, we performed targeted measurement of key intermediate metabolites of PPP. We detected a significantly higher relative amounts of metabolites in male flies (Figure 5D), which suggests a higher metabolic flux through the PPP in males. Consistent with this, NADPH – a key product of PPP was more abundant in males than in females (Figure 5E). Given the prominent role of PPP in the antioxidant defense^81–83^, we hypothesized that elevated PPP activity in males contributes to their resistance to oxidative stress and, thus, to *P. entomophila* infection. To test this hypothesis, we used double mutant *Pgd Zw*. This mutant is lacking functional glucose 6-phosphate dehydrogenase (*Zwischenferment* or *Zw*), the first enzyme in the oxidative PPP, and 6-phosphogluconate dehydrogenase (*Pgd*), the third enzyme in the oxidative PPP^84^. Survival analysis showed that *Pgd Zw* mutant males were as sensitive as females to *P. entomophila* infection (Figure 5F, 5F’) and exhibited intestinal transit blockage (Figure 5G). Thus, PPP is necessary for males to resist *P. entomophila* infection and to clear the pathogen via intestinal peristalsis.

**Figure 5.**
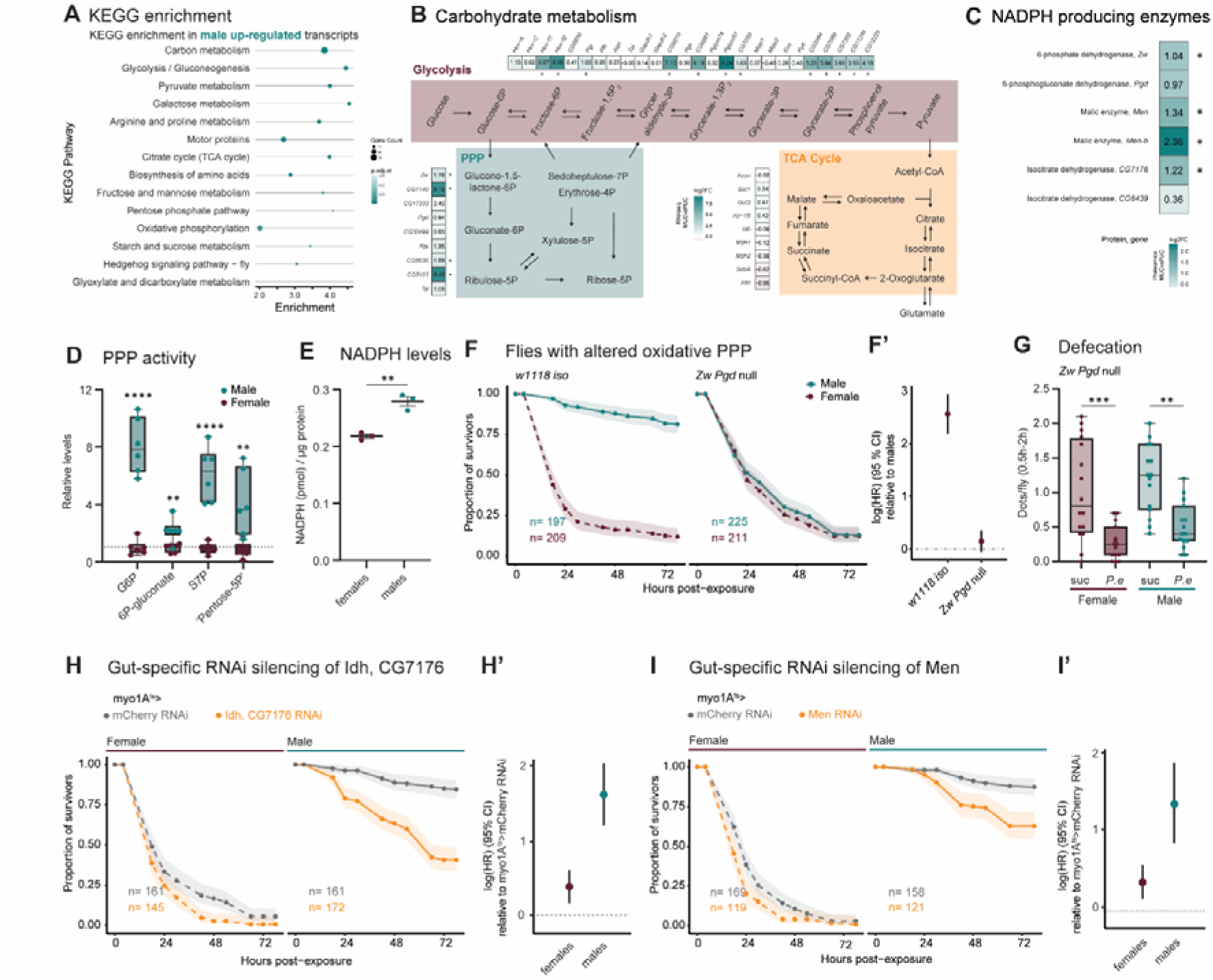
Abolishing differences in NADPH metabolism removes sexual dimorphism in susceptibility to infection. (A) KEGG pathway analysis of genes differentially-regulated between male and female guts under sucrose-fed (control) conditions. (B) The schematic of glycolysis, pentose phosphate pathway (PPP) and tricarboxylic acid (TCA) cycle with respective heatmap showing expression differences (log_2_FC) of selected genes between male and female sucrose-fed (control) guts. Significance (padj < 0.1, |log_2_FC| > 1.5) indicated by *. (C) Heatmap showing abundance difference (log_2_FC) of NADPH producing enzymes between male and female sucrose-fed (control) guts. Significance (padj < 0.05, |log2FC| > 1) indicated by *. (D) Dot plots show metabolite levels normalized to the total protein content of whole-body homogenates relative to mean of female control. “Pentose-5P” represents pentose phosphates; ribose-5-P and ribulose-5-P. Significance by unpaired Student’s t test. (E) Dot plots show the NADPH level of five pooled whole-body homogenates normalized to protein concentration. Mean ± SEM (N=3). Significance by unpaired Student’s t test. **(F and F’)** Survival curves with 95% confidence intervals (shaded area) and hazard ratios with 95% confidence intervals of *w^1118^* iso (control) flies and *Zw Pgd* double null mutant flies (flies lacking two NADPH producing enzymes Zw and Pgd in the oxidative PPP) upon exposure to *P. entomophila*. **(G)** The defecation of female and male *Zw Pgd* double null mutant flies measured 0.5 to 2 h after exposure to blue-dyed sucrose (control) or *P. entomophila* in 0.5 h protocol. Significance by two-way ANOVA with Šídák’s multiple comparisons test. Interaction (Sex x Treatment) p = n.s. **(H – I’)** Survival curves with 95% confidence intervals (shaded area) and hazard ratios with 95% confidence intervals of female and male *myo1A^ts^>mCherry* RNAi (control) and: **(H and H’)** *myo1A^ts^>Idh*, *CG7176* RNAi (flies with knockout of NADPH producing enzyme Idh in the enterocytes), **(I and I’)** *myo1A^ts^>Men* RNAi (flies with knockout of NADPH producing enzyme Men in the enterocytes) upon exposure to *P. entomophila*. For detailed sample sizes and statistical analyses, see Table S9.

Considering that NADPH is a key metabolite of PPP mediating protection against oxidative stress via the production of antioxidants, such as glutathione, we tested the role of additional NADPH-producing enzymes during *P. entomophila* infection. Besides Zw and Pgd, the two enzymes of PPP, the NADPH metabolic network in *Drosophila* includes cytosolic malate dehydrogenase (Malic enzyme, Men), mitochondrial malate dehydrogenase (Malic enzyme b, Men-b), cytosolic isocitrate dehydrogenase (Idh, CG7176), and mitochondrial isocitrate dehydrogenase (Idh, CG6439)^85^. Among these, Zw and Men-b were expressed stronger in male guts when comparing transcripts (Figure 5C), but all of them (except Pgd and mitochondrial Idh, CG6439) had higher protein levels (Figure 5C). Enterocyte-specific knockdown of cytosolic Idh (CG7176) and Men by RNAi resulted in increased susceptibility of males to *P. entomophila* infection (Figure 5H, 5H’, 5I, 5I’). Knockdown of mitochondrial Idh CG6439 had no significant effect on survival (Figure 5D, S5D’). Thus, cytosolic NADPH-producing enzymes are required for increased survival of males to *P. entomophila* infection, likely by providing protection against oxidative stress and consequent suppression of pathogen clearance via gut peristalsis.

### *P. entomophila* virulence regulator Hfq is expressed higher in female guts

Our results suggest that male and female intestines represent distinct environments affecting not only the host defense mechanisms but also potentially the pathogen. To test whether a sexually dimorphic gut environment results in a sex-specific pathogen response, we undertook a proteomics approach to identify proteins differentially produced by *P. entomophila* in the guts of each sex. We detected several *P. entomophila* proteins that differed significantly in abundance between male and female guts (Figure 6A, 6B, listed in Table S5). Among these proteins, RNA chaperon Hfq, which has a role in pathogen virulence in other studies^65–67,86^, was the protein with the largest difference at 6 h (Table S5). We hypothesized that higher expression of Hfq in female flies would result in higher *P. entomophila* virulence, and consequently lower survival of female flies.

**Figure 6.**
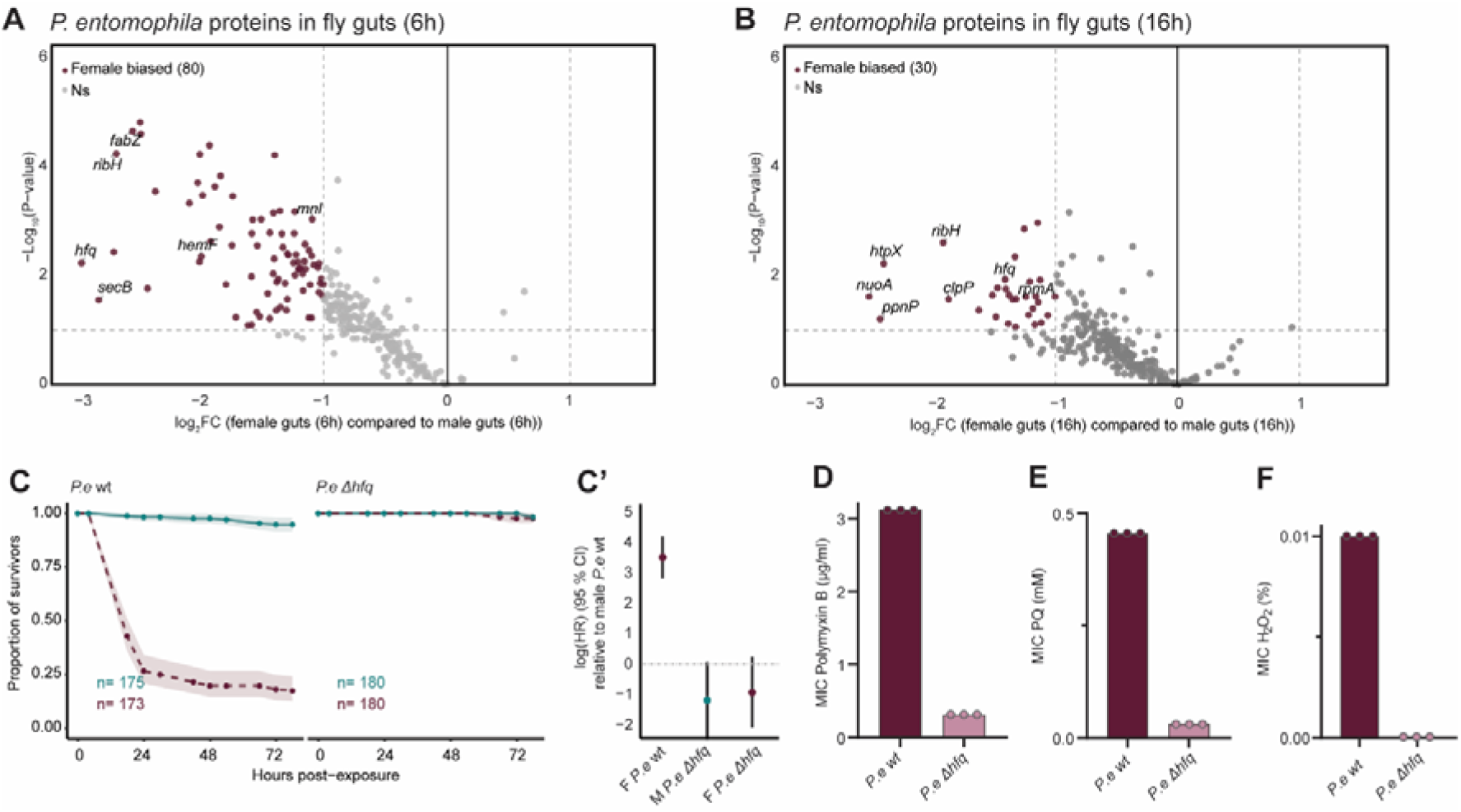
*P. entomophila* virulence regulator Hfq is expressed higher in female guts. **(A and B)** Volcano plots of differentially abundant *P. entomophila* proteins (│log_2_FC│≥ 1 and padj cut-off 0.05) between infected male and female guts **(A)** 6 h and **(B)** 16 h post-exposure to *P. entomophila* (N = 5 independent samples, each with 30 guts pooled). **(C and C’)** Survival curves with 95% confidence intervals (shaded area) and hazard ratios with 95% confidence intervals upon exposure to wild-type *P. entomophila* and *P. entomophila* Δ*hfq*. **(D – F)** Minimum inhibitory concentration (MIC) values of **(D)** polymyxin B, **(E)** paraquat, or (F) hydrogen peroxide against wild-type *P. entomophila* and *P. entomophila* Δ*hfq* mutant (N = 3). For detailed sample sizes and statistical analyses, see Table S9.

To test whether Hfq plays a role in *P. entomophila* virulence in *Drosophila*, we infected flies with the *P. entomophila* Δ*hfq* mutant^87^. Survival analysis showed that the Δ*hfq* mutant was avirulent, and both male and female flies survived the infection (Figure 6C, 6C’). Given the prominent role of Hfq in bacterial stress response^88^, we wondered whether Hfq contributed to the bacterial evasion of female *Drosophila* immune effectors such as ROS and AMPs. To test this, we performed minimum inhibitory concentration (MIC) assay with compounds that should mimic the action of ROS and AMPs^89–92^. The *P. entomophila* Δ*hfq* mutant was more sensitive to all compounds that we tested: polymyxin B (Figure 6D), paraquat (Figure 6E), or hydrogen peroxide (H_2_O_2_) (Figure 6F), suggesting that Hfq is necessary for bacterial ability to counteract host immune effectors. Overall, these findings suggest that the crucial regulator of virulence and stress response Hfq, is expressed higher in the female gut environment, contributing to female-biased virulence of *P. entomophila*.

## DISCUSSION

We report differences between male and female *Drosophila* in susceptibility to intestinal infection. Our results support the following model. Upon ingestion, *P. entomophila* induces excessive oxidative burst in female guts, leading to defecation blockage, pathogen persistence, and death of the flies due to previously reported gut damage. Male flies overcome the pathogen-induced oxidative burst due to elevated basal activity of a key antioxidant system – PPP. This allows males to retain intestinal transit and defecation during infection which expels the pathogens. We further showed that the pathogen produced more of the virulence regulator Hfq in female guts, suggesting that the pathogen behaved differently in male and in female hosts. Our work not only uncovers the mechanistic underpinnings of sexual dimorphism in *Drosophila* susceptibility to intestinal infection but also expands our understanding of virulence strategies utilized by intestinal pathogens.

Our study identified sex differences in the defecation frequency as a key factor explaining the sexual dimorphism in pathogen load and fly survival. Specifically, while defecation was completely blocked in females after *P. entomophila* infection, it was significantly reduced but still retained in males, suggesting that males likely eliminate the pathogen via expulsion. Pathogen clearance via intestinal peristalsis and defecation is a conserved defense mechanism against intestinal infections^93–95^. In *Drosophila*, the following signaling cascade is initiated by bacteria to trigger gut peristalsis. Uracil secreted by ingested pathogens induces DUOX-dependent production of ROS (hypochlorous acid, HOCl) which is released in the gut lumen. HOCl in turn binds to TRPA1 receptor on enteroendocrine cells which secrete large amounts of DH31. DH31 reaches the visceral muscle (VM) where it binds its receptor DH-31R, promoting spasms of the VM to expel bacteria^48,49^. *P. entomophila* is an uracil producer that activates ROS production via DUOX. However in contrast to non-lethal pathogens, like *Pectobacterium carotovorum* (*Ecc15)*, *P. entomophila* induces a disproportionally high level of intestinal ROS^43^. Although prior studies have also shown associations between resistance to *P. entomophila* and intestinal ROS levels, we established a link between the two^60,96^.

While *Ecc15* induces defecation in flies which facilitates pathogen expulsion, our results show that the high level of ROS blocks defecation, hence the bacteria persists in the gut, which kills the flies. The ROS/TrpA1/Dh31 signaling axis is also known to be necessary to trap bacteria. It blocks the pathogenic bacteria in the larvae anterior midgut and kills them by AMPs^47^. However, such trapping strategy is unlikely to be successful against *P. entomophila,* since it blocks translation and production of AMPs, the host would not be able to eliminate the pathogen. Thus, whereas low level of ROS is necessary to trigger gut peristalsis and pathogen clearance, excessive ROS does the opposite – it blocks defecation. Hence, ROS levels need to be fine-tuned to avoid damaging side effects while preserving physiological functions, something that *Drosophila* seem unable to do in the context of *P. entomophila* infection, suggesting that their immune response is maladapted to respond to this infection. Greater immune response in females is a conserved across taxa feature^8^, which contributes to lower intensity and prevalence of many infections in females, but it may increase disease symptoms and severity among females compared with males^13^.

To understand the basis of sexual dimorphism in ROS production and oxidative stress resistance, we looked at sex differences in gene and protein expression under uninfected conditions. Consistent with previous studies^61,62^, we found higher expression of carbohydrate metabolism genes, particularly those involved in glycolysis and PPP in male guts. We confirmed elevated activity of PPP in males using metabolomic analysis which identified increased levels of PPP intermediates, specifically NADPH, in males. NADPH is essential for the enzyme glutathione reductase to regenerate reduced glutathione (GSH) from oxidized glutathione (GSSG). GSH is a key antioxidant that protects cells from oxidative stress^97^. Therefore, PPP-generated NADPH likely provides protection against oxidative stress during the infection by increasing the amount of GSH. In agreement with this hypothesis, male *Drosophila* mutants with whole body alteration of PPP died similarly to females when infected and their defecation was also blocked. This indicates that sex differences in NADPH and antioxidant capacity prior to infection mediate observed differences in survival. But what is the cause of sexual dimorphism in carbohydrate metabolism? Hudry et al^62^ identified that interorgan communication between male gonad and gut determines male bias in the carbohydrate metabolism. Specifically, male gonad activates JAK-STAT signaling in the enterocytes of adjacent gut region consequently inducing the expression carbohydrate metabolism genes. Hence, sexual dimorphism in survival to intestinal infection likely stems from sex differences in gut metabolism driven by JAK-STAT pathway-mediated testis-gut interorgan communication.

Considering that male and female guts represent distinct environments, we hypothesized that the pathogen might respond differently to each sex conditions. Using dual proteomics approach, we were able to not only characterize the host response to infection but also to obtain unique insights into the pathogen response to the host environment. Specifically, we detected higher abundance of several *P. entomophila* proteins in female flies. Among these proteins, we further characterized the RNA chaperon Hfq – a known virulence regulator^66^. We showed that *P. entomophila* Δ*hfq* mutant was avirulent to flies and exhibited increased susceptibility to oxidative stress and antimicrobials mimicking the action of *Drosophila* AMPs. Thus, *P. entomophila* Δ*hfq* mutant exhibits typical phenotypes of attenuated virulence and susceptibility to host defense mechanisms previously reported for *hfq* mutants in other bacteria^65,67,88^. Since we detected decreased abundance of Hfq protein in male guts, this likely leads to decreased pathogen virulence and ability to resist host immune effectors further contributing to the survival of male flies. Our study uncovered an example when gut metabolic environment not only affects the host defenses against the pathogen but also modulates the pathogen virulence. However, the mechanistic basis of Hfq’s differential expression between male and female guts remain to be discovered. Given a prominent role of Hfq is bacterial stress response^88^, it is possible that excessive oxidative stress that occurs specifically in female guts leads to elevated Hfq production as part of *P. entomophila* stress response, consequently increasing pathogen virulence and resistance to host immune effectors. This possibility remains to be tested. Also, considering the pleiotropic role of Hfq in bacterial physiology^66^, further experiments are needed to identify the exact Hfq-regulated processes contributing to virulence.

Besides Hfq, the other *P. entomophila* proteins that are expressed higher in female guts, might also contribute to heightened pathogen virulence to females. For instance, pore-forming toxin monalysin was more abundant in female guts at 6 h post infection. Monalysin is a described virulence factor of *P. entomophila* which participates in the damage to intestinal cells^57^. *P. entomophila-*induced translation blockage and consequent inhibition of immune response and tissue repair were also attributed to the monalysin^43^. Considering a prominent role of pore-forming toxins in the inhibition of gut peristalsis^98^, monalysin is a likely candidate contributing to defecation blockage and *P. entomophila* persistence in female flies.

Our discovery of the sex dimorphic response of the pathogen to the host environment has important implications for pathogen virulence. Given that male and female intestines represent distinct environments, the pathogen experiences different selection pressures in these environments which could lead to divergent evolutionary trajectories. Understanding such long-term consequences of sexual dimorphism in response to infection for pathogen evolution would be an important avenue for future studies.

## STAR+METHODS

### Arrive protocol

The ARRIVE 2.0 guidelines were followed in planning, conducting and reporting *in vivo* experiments^99^.Data used to prepare graphs are summarized in Table S6.

### Drosophila husbandry

*Drosophila* stocks were raised in a light:dark cycle (12 h:12 h) at 25°C, on a standard cornmeal/agar medium (6.2 g agar, 58.8 g cornmeal, 58.8 g inactivated dried yeast, 26.5 ml of a 10% solution of methyl-paraben in 85% ethanol, 60 ml fruit juice, 4.8 ml 99% propionic acid for 1 L). For maintaining flies, flies were transferred to fresh vials every 2-3 days, and fly density was kept to a maximum of 15 flies per vial. At day 2 after eclosion, all emerged adults were transferred in new vials where they were left to mate for 2-3 days. Flies were then sorted using CO_2_ in vials with 20 flies per sex the evening before experiment, unless co-housing the sexes was the aim of the experiment. Flies were then kept at 29°C, unless otherwise stated. Axenic flies were generated by egg bleaching as previously described^91,100^.

### *Drosophila* strains and crosses

*Drosophila* strains used in this study are listed in Table S7. For preparing crosses, five males carrying UAS transgenes were placed together with five virgin females carrying GAL4 driver and transferred to fresh vials every 2-3 days at 18°C. At day 2 after eclosion, all emerged adults were transferred to vials and kept for 2 days at 18°C to allow mating, and then transferred to 29°C where they were kept for at least 2 days before sorting for experiments.

### Oral infection and survival assay

Unless otherwise indicated, the previously established protocol (Figure 1A) was used for the oral infection of flies^43^. *Pseudomonas entomophila* L48 strain was obtained from Bruno Lemaitre’s lab. Previously published *P. entomophila* Δ*hfq* mutant^87^ was kindly provided by Dr. Edna Bode (Max Planck Institute for Terrestrial Microbiology)*. P. entomophila* strains utilized in this study were grown directly from frozen 50% glycerol stocks. Briefly, 10 µL of the stocks was incubated in 20 mL of LB medium in a 50 mL flask overnight (∼16 h) at 29°C with shaking at 175 rpm. The following day, the overnight culture was diluted 1:16 in 150 mL of LB medium and incubated in 500 mL flasks under the same conditions for a minimum of an additional 24 hours. Before the infection experiment, the cultures were centrifuged at 3500 g and 4°C for 15 minutes, and their concentration was estimated to the optical density (OD) of 200 using PBS. The bacterial suspension was then mixed 1:1 with a 5% sucrose solution prior to the infection experiment, and 150 µL of this mixture was pipetted into vials containing standard food and filter papers (referred to as infection vials). Before transferring mated flies (20 per vial) to the infection vials, the flies were kept in empty vials for 2 hours at 29°C (dry starvation). Flies were always infected at 6 Zeitgeber Time (the middle of the 12-hour light cycle) and were kept in the infection vials for 24 hours (unless 0.5 h protocol was used where flies were kept in the infection vials for 0.5 hours) before being transferred to conventional fly food vials. All infections were performed at 29°C. Flies were counted at the following time points: 0.5 h, 18 h, 24 h, 30 h, 42 h, 48 h, 54 h, 66 h, 72 h, and 78 h. Flies counted as ’dead’ at the 0.5 h mark were excluded from the survival analysis due to mortality caused by drowning. All experiments were performed at least twice on different days with at least two vials per day, each containing around 20 flies.

### GWAS (Survival, Analysis, and Validation)

- **a) Survival**

Survival of DGRP lines was performed as above with addition of ‘stricter’ control of mating density (vials where 12 males and 12 females were mated for 48 hours before sorting the flies for experiments) and additionally, counting of the dead flies was performed every 3 h instead of every 6 h during 12 h light cycle.

**- (b) Analysis**

We performed three GWA analyses for each data set: males dataset (HR of male DGRP line relative to the reference line male w1118 iso), females dataset HR of female DGRP line relative to the reference line female w1118 iso), and sexual dimorphism dataset. (HR of female relative to the male of the DGRP line). HR was estimated using a Cox proportional hazards mode (coxph(Surv(Time_to_death_in_day,Censor) ∼ DGRP_lines + DGRP_lines:Sex + strata(Day_of_exposure)). This allowed to calculate the hazard ratios (HR) for each genotype relative to w1118iso line, which was present in every experiment. To identify SNPs associated with variation in HR, we performed a GWAS using a linear model (lm(HR ∼ SNP, weights = 1/(SE^2))). Weighting by the inverse of the variance (1/SE²) allows to give less weight to genotypes with badly estimated HR in the linear model.

- **(c) Functional validation**

To obtain an unbiased approach in gene selection, genes were selected by three authors (D.D., I.I., M.R.). We used multiple criteria such as the previously described function, the functional impact of the mutation as predicted by Ensembl Variant Effect Predictor (i.e. missense, UTR, within gene or intron/splice region), our transcriptomic and proteomics data, and the size of the effects observed on the DGRP lines (i.e. log (HR)). Validation of genes was performed by performing three independent survival assays (predetermined, crosses preparation on separate days with different flies) on crosses where selected candidate genes was downregulated by RNAi in using enterocyte specific GAL4 driver. For preparation of crosses (described above), all GAL4 virgins were collected over a period of five days and then on the day of cross preparation mixed altogether and randomly separated into vials with appropriate males. Survival assays were done in parallel (all genes) and counting of dead flies was performed every 3 hours during 12 h light cycle.

### Food intake quantification

Quantifying the amount of ingested food was done by including blue dye (Food Blue No. 1, TCI) in infection mixture as described previously^48^. *P. entomophila* was mixed in a 1:1 ratio with a 5% sucrose solution containing 1% blue dye. Feeding was interrupted after 0.5 h, and five flies were immediately transferred to pre-weighed tubes containing glass beads and 500 μL of 1x PBS with 0.1% Triton X-100 (PBST). The tubes with the flies were then weighed, allowing for the calculation of the flies’ weight. The samples were then homogenized using a Precellys homogenizer (three cycles of 30 seconds at 7200 rpm) and centrifuged twice (2 mins, maximum speed). 250 μL of supernatant was then transferred to cuvette containing 750 μL of water. Blue dye levels were quantified by measuring absorbance at 630 nm using Ultrospec 2100 pro UV/Vis Spectrometer (Amersham Biosciences). Flies fed with *P. entomophila* mixed 1:1 with 5% sucrose without blue dye were used as a background control. Values were converted to the amount of blue dye (in ng) using a calibration curve obtained from the serial dilutions of a blue dye in PBST and then normalized per weight (in mg) of flies.

### Gut transcriptome comparison (RNA Extraction, RNAseq and GO analysis)

Total RNA was extracted from 30 guts per sample using TRIzol reagent (Invitrogen). RNA quantity and quality were measured using a NanoDrop 2000 (Thermo Scientific). For each condition, RNAs were prepared from three biological replicates on different days. RNA-seq was carried out following standard Illumina protocols, by Novogene (Cambridge, UK). Briefly, RNA quantity, integrity and purity were determined using the Agilent 5400 Fragment Analyzer System (Agilent Technologies). mRNAs were purified from total RNA using poly-T oligonucleotide-attached magnetic beads. After fragmentation, the first-strand cDNA was synthesized using random hexamer primers. Then the second-strand cDNA was synthesized using dUTP, instead of dTTP. The directional library was ready after end repair, A-tailing, adapter ligation, size selection, USER enzyme digestion, amplification and purification. The libraries were quantified with Qubit and checked with bioanalyser for size distribution. Quantified libraries were pooled and sequenced on the Illumina NovaSeq 6000 platform (2u×u150ubp) and generated about 6 Gb of raw data per sample.

We characterized male and female transcriptional response to infection (infected guts compared to sucrose-fed (control) guts of matching sex) and difference between male and female transcriptome under non-infected conditions (male sucrose-fed guts compared to female sucrose-fed guts). Differential expression analysis was done using DESeq2^101^. Significance cut-offs were padj < 0.1, |log_2_FC| > 1.5. Scatter plot with gene categories (genes where the response (infected vs. sucrose-fed (control)) was upregulated or downregulated in both sexes, ‘female biased’ - genes where response (either up or down) was significant only in female flies, and not male flies, ‘male biased’ - genes where response (either up or down) was significant only in male flies, and not female flies) was used to visualize comparison between male and female transcriptional response to infection. KEGG enrichment analysis for gene group lists (comparison of non-infected conditions) was done using enrichKEGG function from the clusterProfiler package. The p-value cutoff was 0.05. The R package ggplot2was used for data visualization.

All raw RNA sequencing data files are available from the SRA database (accession number PRJNA1256345) and can be accessed at https://www.ncbi.nlm.nih.gov/sra/PRJNA1256345 on the day of publication.

### Protein concentration measurement

Protein concentrations were measured by the Pierce BCA Protein Assay Kit (Thermo Fisher Scientific Pierce™ BCA Protein Assay Kit Catalog Numbers 23225 and 23227) according to the manufacturer’s protocol. Briefly, 10 μL of samples were incubated with 200 μL of working reagent and incubated for 30 min at 37°C. Absorbance measurements (in duplicates) were performed at 562unm using a plate reader Infinite M Plex Microplate Reader (Tecan). For incompatible samples, Pierce™ 660 nm Protein Assay Reagent was used. Briefly, 10 μL of samples were incubated with 150 µL of the working reagent, mixed for 1 minute, incubated at room temperature (RT) for 5 minutes. Absorbance measurements (in duplicates) were performed at 660unm using a plate reader Infinite M Plex Microplate Reader (Tecan).

### Gut Proteome Comparison

At the indicated timepoints, 30 guts were dissected in 150 µl of protein extraction buffer (100 mM Tris-Cl pH 8, 2% SDS, protease inhibitor) on ice. Following incubation (95°C for 2 minutes) and homogenization (6000 rpm for 30 seconds, Precellys homogenizer), samples were centrifuges (max speed, 10 min, 4°C). Supernatant was collected into Eppendorf tubes. Protein measurement was performed as described above (Pierce BCA Protein Assay Kit). Samples were stored at -80°C before being analyzed by High Throughput Mass Spectrometry Core Facility (Charité).

**(a) Sample preparation (SP3)**

Gut protein extracts were processed using the SP3 protocol as previously described with one-step reduction and alkylation^102^.

Briefly, 16.6 μl of reduction and alkylation buffer (40 mM Tris(2-carboxyethyl)phosphine, 160 mM Chloroacetamide, 200 mM Ammonium bicarbonate, 4% SDS) was added, samples were incubated at 95 °C for 5 min and cooled to room temperature. To bind the proteins, 250 μg of paramagnetic beads (1:1 ratio hydrophilic/hydrophobic) were used, and the proteins were precipitated by adding 50% Acetonitrile. Samples were washed twice with 80% EtOH and once with 100% Acetonitrile. 35µl of 100 mM Ammonium bicarbonate and Trypsin/LysC (0.1 µg/µl stock solution protein: enzyme ratio of 1:50 (w/w)) were added and the lysates were incubated at 37°C with shaking overnight. The reaction was stopped by adding formic acid to a final concentration of 0.1%. Peptide concentration was determined using Pierce Quantitative Fluorometric Peptide Assay Kit, and peptide mixtures were analyzed by LC-MS/MS without further conditioning or clean-up.

**(b) Proteome analysis by DIA LC-MS**

Peptide separation was accomplished in a data independent acquisition (DIA) mode for a 63-minute in water to acetonitrile active gradient on an Ultimate 3000 RSLCnano HPLC coupled to a Q-Exactive Plus mass spectrometer (both ThermoFisher Scientific). 1 μg digested peptides were trapped on a column (PepMap C18, 5 mm x 300 μm x 5 μm, 100 Ǻ, Thermo Fisher Scientific) with buffer containing 2:98 (v/v) acetonitrile/water in 0.1% (v/v) trifluoroacetic acid, flow rate of 20 μl/min for 3 min. The peptide mixture was separated on a C18 column (Acclaim PepMap C18, 2 μm; 100 Å; 75μm, Thermo Fisher Scientific) using a linear gradient from 5% to 28% buffer B over 63 minutes, followed by an increase to 95% buffer B in 2 minutes, and a 5-minute wash with 95% buffer B before a 20-minute equilibration with initial conditions. Buffer A is made of 0.1% formic acid in LC-MS-grade water; buffer B is made of 80% acetonitrile and 0.1% formic acid mixed with LC-MS-grade water. Total acquisition time was 100 min. The Orbitrap worked in centroid mode including a duty cycle consisted of one MS1 scan at 70,000 resolution power with maximum injection time 300 ms and 3e6 AGC target followed by 40 variable MS2 scans using an 0.5 da overlapping window pattern. The acquisition started with window length of 25 MS2 scans at 12.5 da; followed by 7 windows with 25 da, then the last 8 windows were set to 62.5 da. The resolution of precursor MS spectra (m/z 378-1370) was set to 17,500 after accumulating ions for 110 ms to achieve a target value of 3e6. Mass spectrometric settings were set to: spray voltage, 2.0 kV; no sheath and auxiliary gas flow; heated capillary temperature, 275 °C; normalized HCD collision energy 27%. Additionally, the background ions m/z 391.2843 and 445.1200 acted as lock mass.

**(c) Data analysis**

Raw data were processed using DIA-NN 1.8.1^103^ with MS2 and MS1 mass accuracies set to 20 and 10 ppm, respectively. The output was filtered at 1% FDR on peptide level.

Quantification strategy was “Robust LC with high precision”. A spectral library free search with activated match between runs (MBR) was used^104^.

We characterized male and female proteomic response to infection (infected guts compared to sucrose-fed (control) guts of matching sex) and difference between male and female proteome under non-infected conditions (male sucrose-fed guts compared to female sucrose-fed guts). The differential analysis of quantitative proteomics data was performed in Perseus v2.0.6.0. Significance cut-offs were padj < 0.05, |log2FC| > 1. Scatter plot with protein categories (proteins where the response (infected vs. sucrose-fed (control)) was upregulated or downregulated in both sexes, ‘female biased’ - proteins where response (either up or down) was significant only in female flies, and not male flies, ‘male biased’ - proteins where response (either up or down) was significant only in male flies, and not female flies) was used to visualize comparison between male and female proteomic response to infection. KEGG enrichment analysis for protein group lists (comparison of non-infected conditions) was done using enrichKEGG function from the clusterProfiler package. The p-value cutoff was 0.05. The R packages ggplot2 was used for data visualization.

The mass spectrometry proteomics data have been deposited to the ProteomeXchange Consortium via the PRIDE partner repository with the dataset identifier PXD064190.

### Bacterial load measurement

Bacteria loads were recorded by CFUs after infecting flies for 0.5 hours and then flipping them to conventional vials. Samples from pooled groups of five flies were collected at 0.5, 2, 4, 6, and 24 hours since initial exposure to *P. entomophila*. The flies were first surface sterilized in 70% ethanol for 1 minute, following three washes in sterile PBS for 30 sec, and then homogenized in 500 μL of sterile PBS for 30 seconds at 6000 rpm using a Precellys 24 instrument (Bertin Technologies, France). Serial 10-fold dilutions, ranging from 10⁻^1^ to 10⁻^6^, were made and plated on LB plates using an automatic diluter and plater (easySpiral Dilute (Interscience, France)). After overnight incubation (proxy 18 h) at 30°C, colonies were counted with an automatic colony counter (Scan 1200 (Interscience, France)), along with its accompanying software. As a control, flies fed with sucrose were processed in parallel with infected flies. Since the control plates were negative for bacterial colonies, it was assumed that all colonies that grew at plates of infected flies are *P. entomophila*.

### Measuring defecation in adult flies

To measure defecation, we prepared and infected flies with a suspension containing blue dye as described above (see: Food intake quantification). Flies were kept on blue suspension for 0.5 h and then transferred to the vial containing conventional non-colored food for an additional 1.5 hours (time points were chosen to reflect bacterial load experiment (see: Figure 3A) where we observed first reduction of bacteria in male flies). Defecation rate was measured by counting ‘defecation dots’ left dried on the inner wall of vials and normalized per number of flies in vial. Experiments were performed at least twice on different days, with a total of at least 10 independent vials per condition, each containing 10 or 20 flies.

### Chemical manipulations

For experiments where the effect of a compound on survival or defecation was tested, experiments were performed and described before, with the exception that either PBS (control) or *P. entomophila* were mixed 1:1 with sucrose-containing compounds. The final concentration (after 1:1 mix) were as follows: N-Methylmaleimide (NMM,Thermo Scientific, Cat# 127080050) was 1 mM, N-Acetyl-L-Cysteine (NAC, Sigma-Aldrich, Cat# A7250) was 20 mM, Methyl Viologen hydrate (98% (paraquat, (Thermo Scientific, Cat# 227320050)) was 20 mM).

### ROS measurement

We used the fluorescent dye 2′,7′-dichlorofluorescein diacetate (H_2_DCF-DA, ≥97%, Invitrogen™, Cat# D399) as an indicator of general oxidative stress^79^. At specified time points, 30 guts were dissected directly into 150 µL of PBS kept on ice. The samples were homogenized using glass beads (30 seconds, 6000 rpm, Precellys 24 instrument (Bertin Technologies, France)). The entire homogenate was transferred to fresh Eppendorf tubes and centrifuged at maximum speed for 10 minutes at 4°C. 70 μL of the supernatant was transferred to a new tube and used for protein measurement, as described above (Pierce BCA Protein Assay Kit), and for measurement of general oxidative stress. 10 µL of gut homogenate was mixed with 190 µL of a 1x PBS solution containing 50 μM H_2_DCF-DA and incubated for 30 minutes at 37°C in darkness. Fluorescence was measured using an excitation wavelength of 488 nm and an emission wavelength of 520 nm. Auto-oxidation of H_2_DCF-DA served as a control. All samples were measured in duplicates and the levels of ROS were normalized as a percentage of relative DCF fluorescence units per microgram of protein in the sample.

### Metabolites Comparison

Five whole flies per sample were used following the 6 h sucrose-feeding (control that was used for RNAseq and proteomics). To extract the metabolites, flies were homogenized using a Precellys homogenizer (6000 rpm, 30 seconds) in a mixture of 100 µL of 50% methanol and 150 µL of chloroform. Following centrifugation (10,000 g, 20 minutes, 4°C), 50 µL of the upper aqueous layer were transferred into HPLC/GC certified vials (Fisherbrand™) on ice. Protein concentration was measured as described above (Pierce™ 660 nm Protein Assay Reagent). Rest of the samples were kept at -80°C before sending to Core facility for metabolomics and small molecules mass spectrometry (Max Planck Institute for Terrestrial Microbiology, Marburg) for metabolomic analysis.

Quantitative metabolite determination was performed using a LC-MS/MS. The chromatographic separation was performed on an Agilent Infinity II 1290 HPLC system using a SeQuant ZIC-pHILIC column (150 × 2.1 mm, 5 μm particle size, peek coated, Merck) connected to a guard column of similar specificity (20 × 2.1 mm, 5 μm particle size, Phenomoenex) a constant flow rate of 0.1 ml/min with mobile phase A with mobile phase comprised of 10 mM ammonium acetate in water, pH 9, supplemented with medronic acid to a final concentration of 5 μM (A) and 10 mM ammonium acetate in 90:10 acetonitrile to water, pH 9, supplemented with medronic acid to a final concentration of 5 μM (B) at 40°C .

The injection volume was 1 µl. The mobile phase profile consisted of the following steps and linear gradients: 0 – 1 min constant at 75% B; 1 – 6 min from 75 to 40% B; 6 to 9 min constant at 40% B; 9 – 9.1 min from 40 to 75% B; 9.1 to 20 min constant at 75% B. An Agilent 6495 ion funnel mass spectrometer was used in positive and negative mode with an electrospray ionization source and the following conditions: ESI spray voltage 2000 V(-)/3500 V(+), nozzle voltage 1000 V, sheath gas 300° C at 20 l/min, nebulizer pressure 20 psig and drying gas 100°C at 11 l/min. Compounds were identified based on their mass transition and retention time compared to standards. Chromatograms were integrated using MassHunter software (Agilent, Santa Clara, CA, USA).

Mass transitions, collision energies, Cell accelerator voltages, and Dwell times have been optimized using chemically pure standards. The parameter settings of all targets are given in Table S8.

## NADPH/NADP+ KIT

The NADP+/NADPH ratio was measured from whole fly lysates using the NADP+/NADPH Quantification Colorimetric Kit (Abcam, Cat# ab65349) according to the manufacturer’s protocol. Colorimetric measurements were performed at 450unm using a plate reader Infinite M Plex Microplate Reader (Tecan). Five whole flies per sample were used following the 6 h sucrose-feeding (control that was used for RNAseq and proteomics). Values were normalized per protein concentration measured as described above (Pierce™ 660 nm Protein Assay Reagent).

## MIC

Overnight bacterial cultures were adjusted to OD 0.1, diluted 1:100, and pipetted (50 µl) into wells of 96 well plates prefilled with ranging dilutions of 50 µl of Polymyxin B (Fischer Scientific) (nine 1:1 dilutions from initial 100 µl/mL and 40 µl/mL solutions (final minimal concertation tested 0.0781 µl/mL Polymyxin)., paraquat (Methyl Viologen hydrate, 98%, thermo scientific) (nine 1:1 dilutions from initial 15 mg/mL and 6 mg/mL solutions (final minimal concertation tested 0.0117 µg/mL paraquat)) or H_2_O_2_ (Hydrogen Peroxide Solution, Sigma-Aldrich) (nine 1:1 dilutions from initial 3% and 1.2% solutions (final minimal concertation tested 0.002 % µg/mL H_2_O_2_). After overnight incubation, the bacterial growth was recorded to determine MIC value. OD600 was measured. Reads were performed in Infinite M Plex Microplate Reader (Tecan).

### Quantification and statistical analysis

Statistical parameters and tests are shown in the figure legends and Table S9. No formal randomization method was used, but to reduce the potential bias, we varied the order of sorting flies, infection vials, sampling and well plate location of samples across different biological replicates. Additionally, control and treatment groups were processed in parallel, and sample sizes were balanced. Blinding was not implemented in this study. No data were excluded from analysis. Data analyses were performed using GraphPad Prism 10 software and R v4.3.2. Kinetics of survival data were shown using cumulative data, and survival analysis was carried with Cox proportional hazards model using Survival R package. Log Hazard ratio are represented to provide the estimate of difference in survival considering all the parameters, including day of the experiment. 95% confidence intervals are provided by the package Survival and are approximated by calculating 1.96 times the standard error. Data visualization was performed with the R packages ggplot2, dplyr, and tidyverse. Statistical significance was determined using either the unpaired Student’s t-test or Mixed-effects model (REML) or one-way or two-way ANOVA, followed by post-hoc tests for multiple comparisons (indicated in the figure legends). Significance was set at p < 0.05, with data presented as mean ± SEM. Significance: *p < 0.05, **p < 0.01, ***p < 0.001, ****p < 0.0001.

## Acknowledgments

We are grateful to Bruno Lemaitre and the Bloomington Drosophila Stock Center (NIH P40OD018537) for fly stocks. We thank Edna Bode and Helge Bode for kindly providing *P. entomophila* Δ*hfq* mutant. We thank the Core Facility High Throughput Mass Spectrometry of the Charité for support in acquisition of the proteomics data and Diane Schad for help with preparation of Graphical Abstract. This work was supported by the Max Planck Society. I.I. also acknowledges the funding from the Deutsche Forschungsgemeinschaft (grants IA 81/2-1 and IA81/3-1) and from the Boehringer Ingelheim Foundation.

## Author Contributions

I.I. initiated the study and acquired funding. M.R., A.A.-R, D.D., and I.I. designed the experiments. M.R., A.A.-R, K.A.M., W.K., N.P., and I.I. performed the experiments. M.R., A.A.-R., N.P., K.A., D.D., and I.I. analyzed the data. D.D. and I.I. supervised M.R., M.R. supervised K.A.M. and W.K. M.R. and I.I. wrote the manuscript with input from all authors.

## Supplemental Information

**Supplementary Figure 1.**
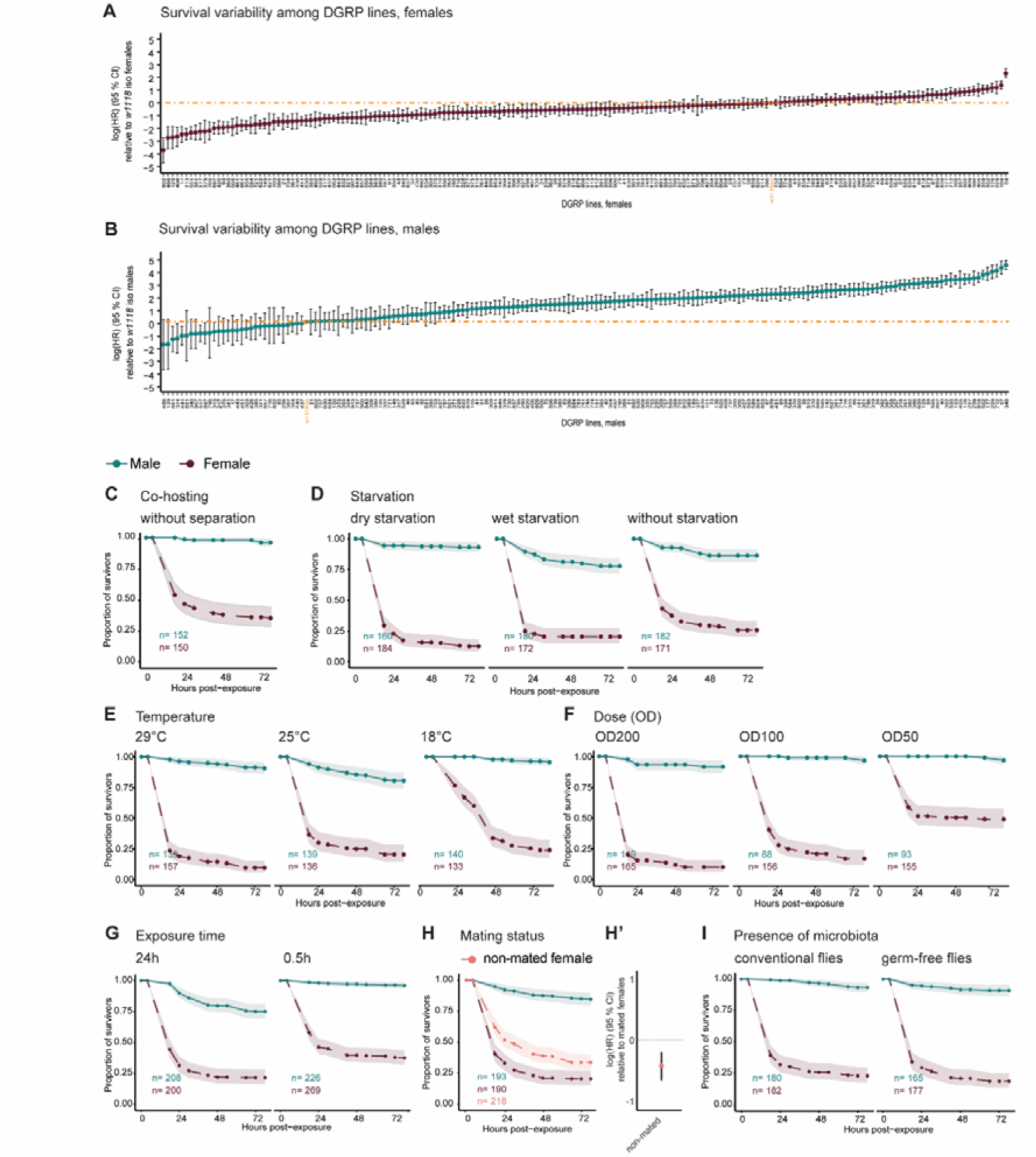
Sexual dimorphism in survival to *P. entomophila* intestinal infection is common across genotype and experimental conditions. **(A and B)** Variability in susceptibility to *P. entomophila* infection across each sex (**(A)** females and **(B)** males) of DGRP lines compared to the appropriate sex of the *w^1118^* iso strain. Data is shown in Figure 1 **C.** **(C – G, H, I)** Survival curves with 95% confidence intervals (shaded area) of *w^1118^* iso upon exposure to *P. entomophila* under different protocol conditions. Data is related to Figure 1 **E – I, K, L**. (H’) Hazard ratios with 95% confidence intervals of non-mated *w^1118^* iso females upon exposure to *P. entomophila* compared to mated *w^1118^*iso females. For detailed sample sizes and statistical analyses, see Table S9.

**Supplementary Figure 2.**
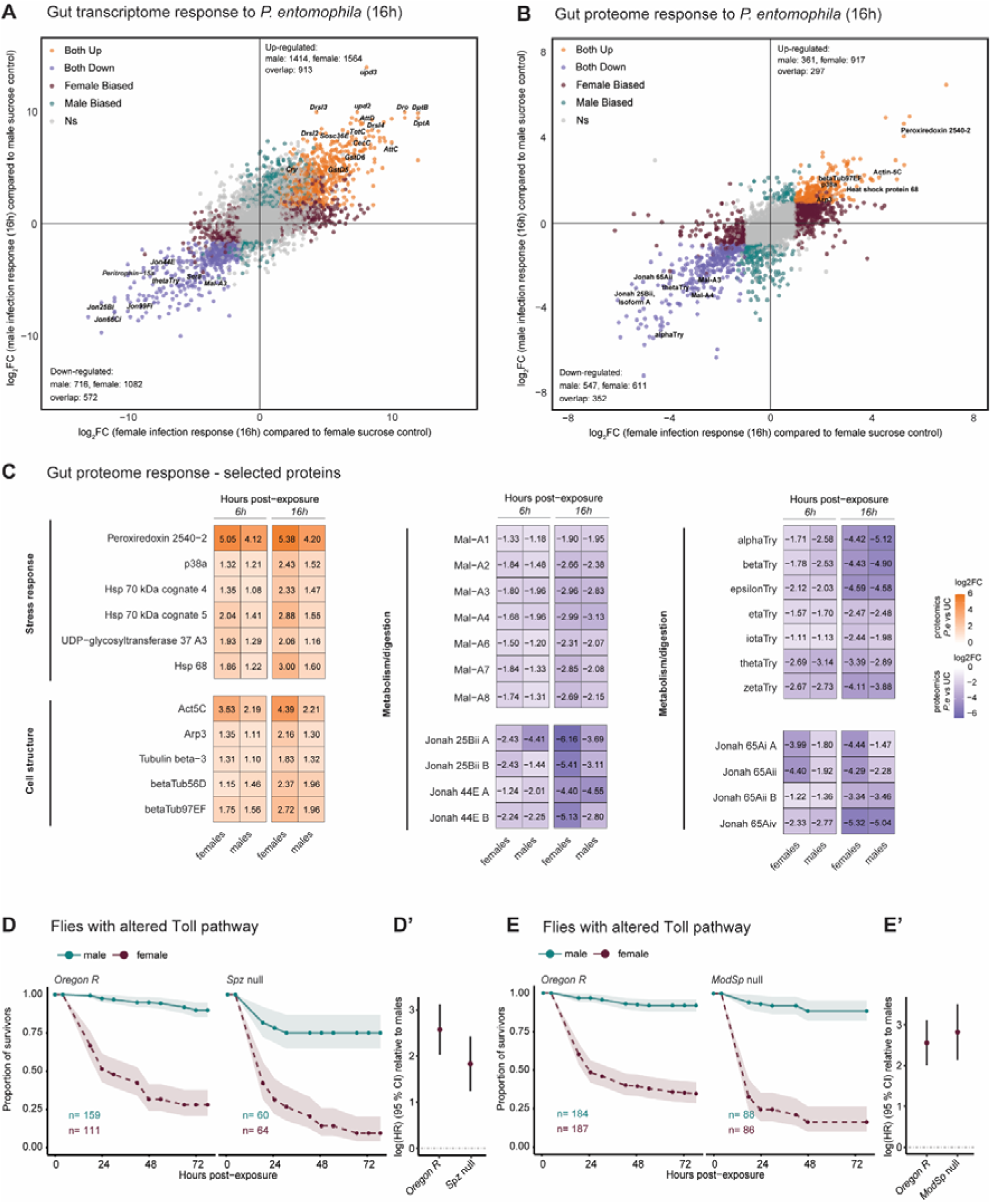
Sexual dimorphism in survival to *P. entomophila* intestinal infection is not explained by the major immune pathways. **(A and B)** Scatter plots representing log_2_FC of **(A)** gene expression or **(B)** protein abundance in female (x) vs male (y) guts 16 h post-exposure to *P. entomophila* (infected guts compared to sucrose-fed (control) guts of matching sex). Significance cut-offs: **(A)** padj < 0.1, |log_2_FC| > 1.5, and **(B)** padj < 0.05, |log_2_FC| > 1. **(A)** (N = 3 independent samples, each with 30 pooled guts). **(B)** (N = 5 independent samples, each with 30 pooled guts). **(C)** Heatmaps showing differences (log_2_FC, infected guts compared to control guts) in the abundance of selected proteins of each sex at 6 h and 16 h post-exposure to *P. entomophila*. **(D and D’)** Survival curves with 95% confidence intervals (shaded area) and hazard ratios with 95% confidence intervals of Oregon R (control) and *spz^rm7^* loss-of-function mutant upon exposure to *P. entomophila*. **(E and E’)** Survival curves with 95% confidence intervals (shaded area) and hazard ratios with 95% confidence intervals of Oregon R (control) and *ModSp* loss-of-function mutant upon exposure to *P. entomophila*. For detailed sample sizes and statistical analyses, see Table S9.

**Supplementary Figure 3.**
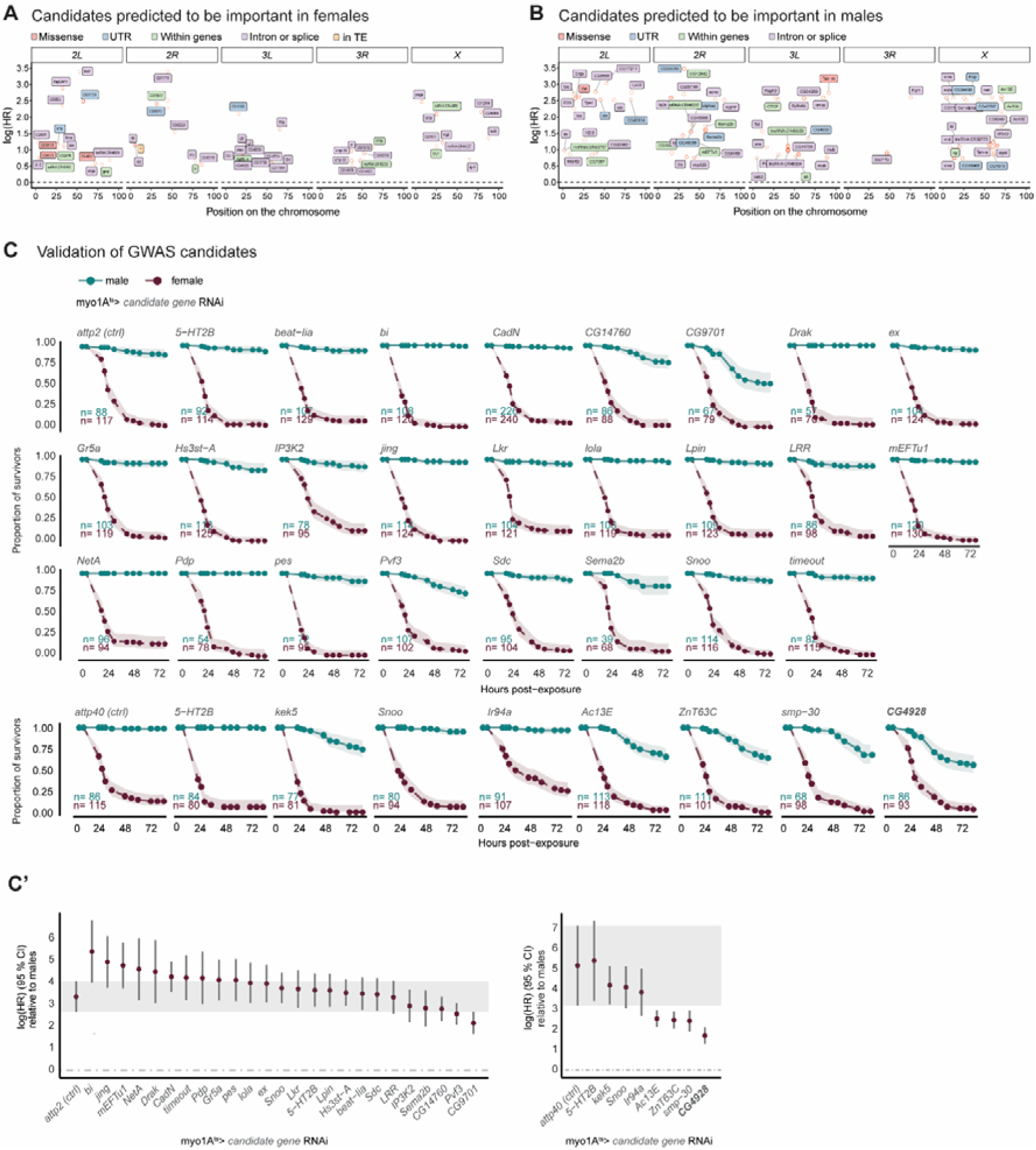
Validation of GWAS-identified candidates revealed new mediators of male survival after infection. **(A and B)** Candidate genes underlying natural variation in the susceptibility to *P. entomophila* gut infection among **(A)** female and **(B)** male flies. **(C and C’)** Survival curves with 95% confidence intervals (shaded area) and hazard ratios with 95% confidence intervals of *myo1A^ts^>* crossed with appropriate control (*attp2* RNAi or *attp40* RNAi) and RNAi line for GWAS gene candidates upon exposure to *P. entomophila*. Part of the data is shown in Figures 3H and **3H’**. For detailed sample sizes and statistical analyses, see Table S9.

**Supplementary Figure 4.**
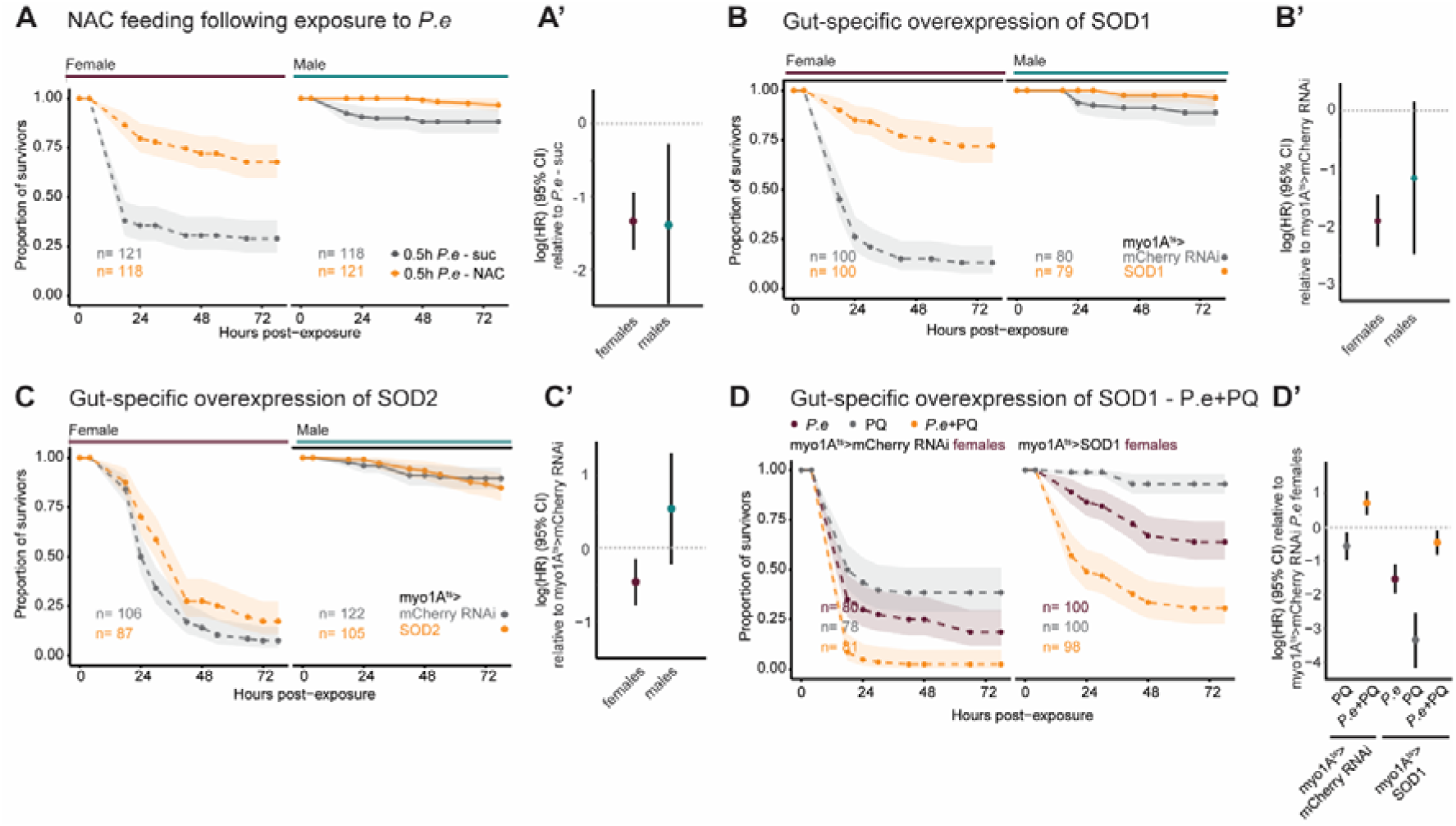
Cytosolic but not mitochondrial oxidative stress determines female susceptibility to infection. **(A and A’)** Survival curves with 95% confidence intervals (shaded area) and hazard ratios with 95% confidence intervals of female and male *w^1118^*iso flies upon exposure to *P. entomophila* (4 h protocol) followed by sucrose (control) or N-Acetyl-L-Cystein (NAC) - supplemented sucrose. **(B and B’)** Survival curves with 95% confidence intervals (shaded area) and hazard ratios with 95% confidence intervals *myo1A^ts^>mCherry* RNAi (control) and *myo1A^ts^>UAS-SOD1* (BL24574) upon exposure to *P. entomophila*. **(C and C’)** Survival curves with 95% confidence intervals (shaded area) and hazard ratios with 95% confidence intervals *myo1A^ts^>mCherry* RNAi (control) *myo1A^ts^>UAS-SOD2* upon exposure to *P. entomophila*. **(D and D’)** Survival curves with 95% confidence intervals (shaded area) and hazard ratios with 95% confidence intervals of female *myo1A^ts^>mCherry* RNAi (control) and *myo1A^ts^>UAS-SOD1* (BL24750) flies upon exposure to *P. entomophila*, paraquat, and *P. entomophila*/paraquat mixture. For detailed sample sizes and statistical analyses, see Table S9.

**Supplementary Figure 5.**
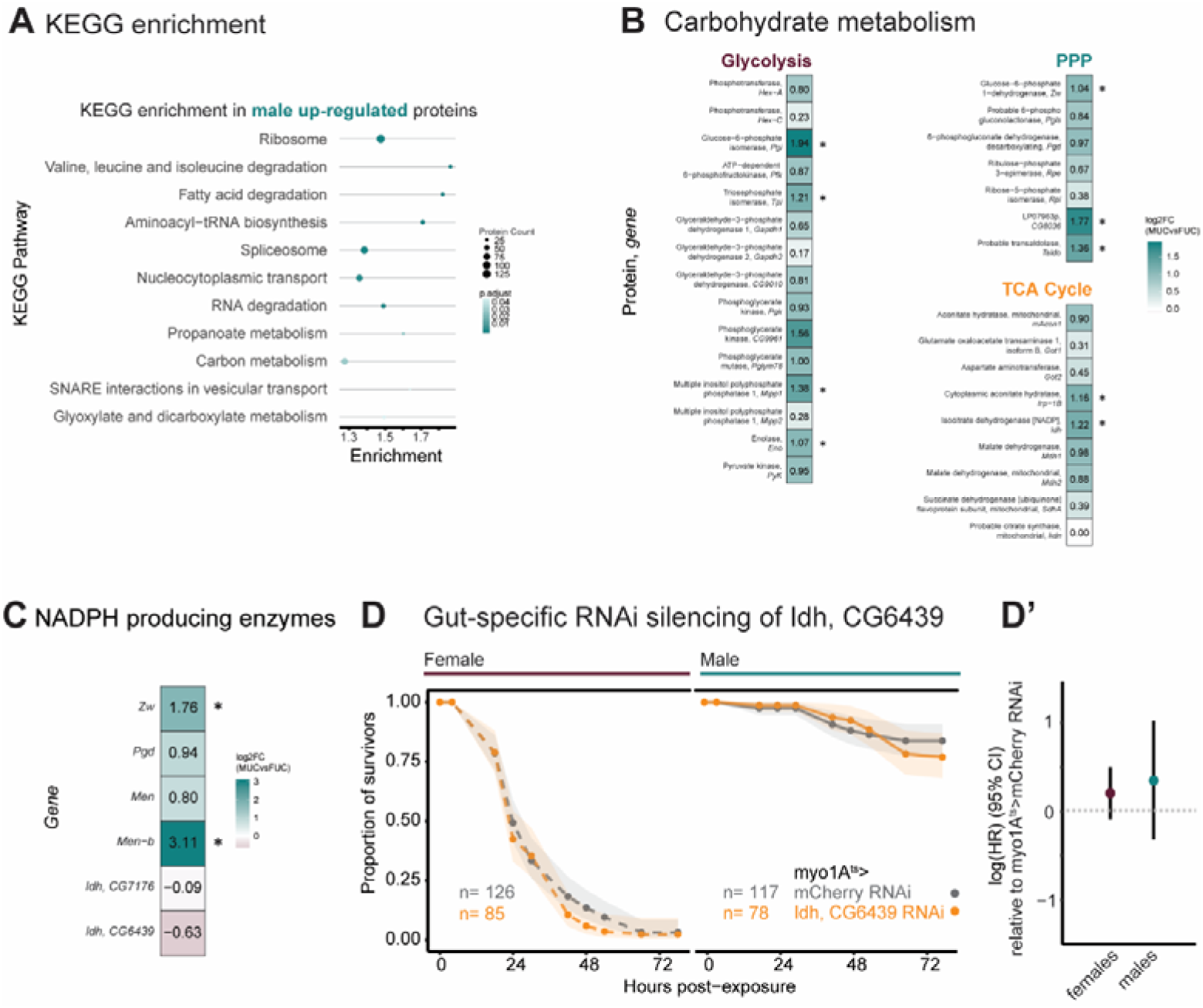
Male bias in NADPH contributes to gut antioxidant capacity needed for survival. **(A)** KEGG pathway analysis of proteins differentially-regulated between male and female guts under sucrose-fed (control) conditions. **(B)** Heatmap showing abundance difference (log_2_FC) of proteins involved in glycolysis, PPP, and TCA cycle between male and female guts under sucrose-fed (control) conditions. Significance (padj < 0.05, |log2FC| > 1) indicated by *. **(C)** Heatmap showing expression level difference (log_2_FC) of NADPH producing enzymes between male and female guts under sucrose-fed (control) conditions. Significance (padj < 0.1, |log2FC| > 1.5) indicated by *. **(D and D’)** Survival curves with 95% confidence intervals (shaded area) and hazard ratios with 95% confidence intervals of female and male *myo1A^ts^>mCherry* RNAi (control) and *myo1A^ts^ >Idh, CG6439* RNAi upon exposure to *P. entomophila*. For detailed sample sizes and statistical analyses, see Table S9.

Table S1. Excel tables listing genes differentially-expressed under infection conditions.

Table S2. Excel tables listing proteins differentially-expressed under infection conditions.

Table S3. Excel tables listing GWAS-identified SNPs and candidate genes.

Table S4. Excel tables listing transcripts and proteins differentially-expressed between male and female guts under basal conditions.

Table S5. Excel tables listing *P. entomophila* proteins with differential abundance between male and female guts.

Table S6. Excel tables with data used to generate graphs.

Table S7. Excel table with the list of fly lines used in this study.

Table S8. Excel table with the parameter settings of target metabolites measured in this study.

Table S9. Excel tables reporting the details of statistical analysis.

